# Adaptive Pathfinding by Nucleokinesis during Amoeboid Migration

**DOI:** 10.1101/2023.05.19.540965

**Authors:** Janina Kroll, Robert Hauschild, Kasia Stefanowski, Monika D. Hermann, Jack Merrin, Lubuna Shafeek, Annette Müller-Taubenberger, Jörg Renkawitz

## Abstract

Motile cells moving in multicellular organisms encounter microenvironments of locally heterogeneous mechanochemical composition. Individual compositional parameters like chemotactic signals, adhesiveness, and pore sizes are well known to be sensed by motile cells, providing individual guidance cues for cellular pathfinding. However, motile cells encounter diverse mechanochemical signals at the same time, raising the question of how cells respond to locally diverse and potentially competing signals on their migration routes. Here, we reveal that motile amoeboid cells require nuclear repositioning, termed nucleokinesis, for adaptive pathfinding in heterogeneous mechanochemical microenvironments. Using mammalian immune cells and the amoeba *Dictyostelium discoideum*, we discover that frequent, rapid and long-distance nucleokinesis is a basic component of amoeboid pathfinding, enabling cells to reorientate quickly between locally competing cues. Amoeboid nucleokinesis comprises a two-step cell polarity switch and is driven by myosin II-forces, sliding the nucleus from a ‘losing’ to the ‘winning’ leading edge to re-adjust the nuclear to the cellular path. Impaired nucleokinesis distorts fast path adaptions and causes cellular arrest in the microenvironment. Our findings establish that nucleokinesis is required for amoeboid cell navigation. Given that motile single-cell amoebae, many immune cells, and some cancer cells utilize an amoeboid migration strategy, these results suggest that amoeboid nucleokinesis underlies cellular navigation during unicellular biology, immunity, and disease.

## Introduction

The ability of cells to navigate their path while moving themselves forward is critical for innate and adaptive immune responses, organismal development, tissue maintenance, and single-cell organisms. During navigational migration, cells are surrounded by local environments that are composed of extracellular matrix, interstitial fluid, and tissue cells in the case of cellular migration in multicellular organisms, or natural environments like soil in the case of motile single-cell organisms. To efficiently navigate through these environments, migrating cells are equipped with mechanisms to follow diverse chemical as well as mechanical cues (Yamada & Sixt, 2019; Moreau *et al*, 2018; Helvert *et al*, 2018; Charras & Sahai, 2014). Most well-investigated are chemotactic signals like chemokines, which are detected by the respective cellular receptors and align intracellular force generation by the actin cytoskeleton towards the chemotactic cue, such as during the recruitment of immune cells to sites of inflammation or into lymphatics (Worbs *et al*, 2017; Hauser *et al*, 2016; Nourshargh & Alon, 2014). Yet at the same time, motile cells also encounter mechanical guidance cues along their paths, such as differently sized pores in the extracellular matrix (Wolf *et al*, 2013; Renkawitz *et al*, 2019).

Navigating motile cells frequently generate new protrusions (‘leading edges’; ‘cell fronts’) next to already existing ones, followed by favoring one of both protrusions along which the path of migration is subsequently orientated. Given that these morphological characteristics are particularly frequent in more shallow gradients (Kay *et al*, 2008; Andrew & Insall, 2007; Swanson & Taylor, 1982; Gerisch & Keller, 1981) and that deficiency of the Arp2/3 regulator Hem1 causes not only a shape simplification into less ramified shapes but also impaired migration along chemokine gradients (Leithner *et al*, 2016), it is likely that these alternative protrusions simultaneously measure guidance cues at alternative local sites in the microenvironment. These explorative protrusions are most evident in fast-migrating amoeboid cells, such as neutrophils and dendritic cells, generating highly dynamic and ramified cell shapes (Fritz-Laylin *et al*, 2017; Renkawitz *et al*, 2019; Driscoll *et al*, 2019). While some principles of how migrating cells coordinate these complex cell shapes have been identified (Hadjitheodorou *et al*, 2023, 2021; Kopf *et al*, 2020; Devreotes *et al*, 2017), it remains entirely unknown how motile cells deal with locally diverse and potentially competing mechanochemical inputs, and how they reorganize their intracellular content along the dominant protrusion once a path has been selected. Here, we address these open questions by investigating how amoeboid cells navigate their path in locally heterogeneous microenvironments.

## Results

### Rapid and long-distance nucleokinesis during amoeboid immune cell migration

As a model system for cell migration in microenvironments of heterogeneous mechanochemical composition, we imaged dendritic cells migrating along a chemotactic gradient (CCL19) while being embedded in a three-dimensional collagen matrix composed of varying pore sizes (Driscoll *et al*, 2019; Renkawitz *et al*, 2019; Schwarz *et al*, 2017; Wolf *et al*, 2013). Migrating dendritic cells frequently showed a ramified cell shape with multiple protrusions, continuously generating new protrusions next to existing ones and eventually favoring one protrusion to select the preferred path of migration (Fig 1A and B, and Movie EV1). In line with the function of the nucleus to act as a mechanical ruler to guide migration along the path of larger pore sizes (Venturini *et al*, 2020; Lomakin *et al*, 2020; Renkawitz *et al*, 2019), the cellular path was frequently identical with the nuclear path (Fig 1B). Yet, in one out of four path decisions we observed the mispositioning of the nucleus into future ‘losing’ protrusions (Fig 1B-E). This deviation of the nuclear path from the cellular path caused a subsequent long-distance repositioning of the nucleus into the winning protrusion, readjusting the nuclear to the cellular path (Movie EV1). Even more often we observed short-range nuclear repositioning events, when the nucleus deformed simultaneously into at least two alternative paths, requiring repositioning of subnuclear parts to the chosen cell path (Fig 1C and Movie EV1).

**Figure 1.**
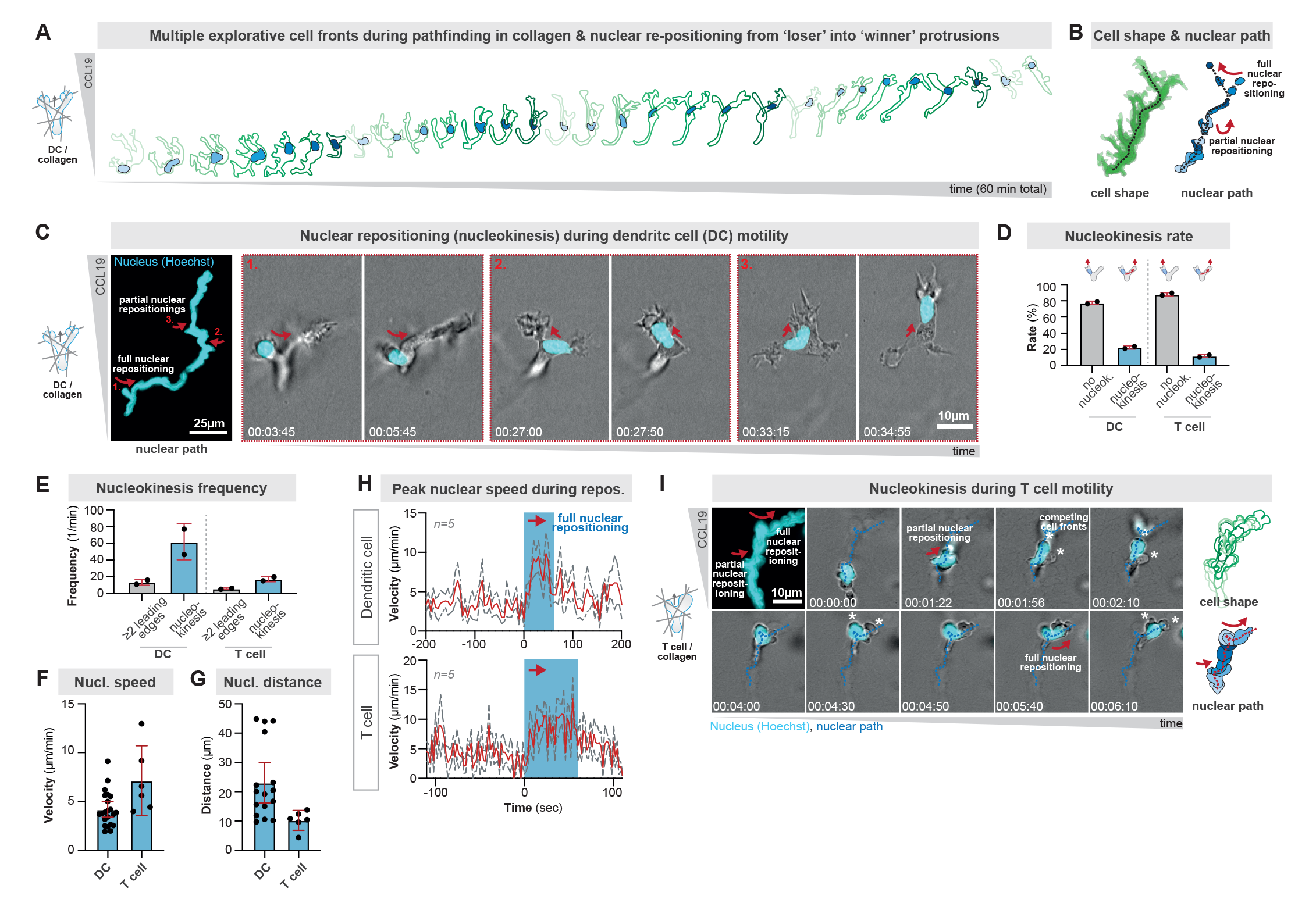
Immune cells employ rapid and long-distance nucleokinesis during amoeboid cell migration. **A.** Cellular outlines (shades of green, time color-coded) of a representative bone marrow-derived mouse dendritic cell (DC) migrating in a three-dimensional collagen matrix along a CCL19 chemokine gradient. The intracellular localization of the DC nucleus during pathfinding is shown in blue (shades of blue, time color-coded). See also ‘Expanded View Movie 1’. **B.** Projection cellular (green) and nuclear (blue) path of the cell in A) over 60 minutes time. Note the two events, in which either large parts of the nucleus or the entire nucleus deviate from the cellular path, required partial and full nuclear repositioning, respectively. **C.** Representative nucleokinesis events in a bone marrow-derived DC migrating through a three-dimensional collagen matrix along a CCL19 chemokine gradient. Highlights in dashed boxes show full (1.) and partial (2. & 3.) nucleokinesis events. Time in hr:min:sec. **D.** Quantification of the rate of full nucleokinesis events in amoeboid DCs and T cells migrating in a collagen matrix: when the migrating cell has a least two major protrusions, quantification of whether the nucleus locates immediately into the dominant protrusion or first locates into the future retracting protrusion, requiring full nucleokinesis into the future dominant protrusion. Data are mean±SD. N=26 DCs (2 replicates, 107 events) and 13 T cells (2 replicates, 49 events). **E.** Quantification of the frequency of full nucleokinesis events in amoeboid DCs and T cells migrating in a collagen matrix: as in D), but showing the frequency of at least two leading edges and full nucleokinesis events per minute. Data are mean±SD. N=26 DCs (2 replicates, 107 events) and 13 T cells (2 replicates, 49 events). **F.** Quantification of the speed of nuclear movement during full nucleokinesis in DCs and T cells. Data are median±95CI. **G.** Quantification of the intracellular distance of nuclear movement during full nucleokinesis in DCs and T cells. Data are median±95CI. **H.** Measurements of nuclear speed before, during, and after full nucleokinesis events in DCs and T cells. The blue box marks the nucleokinesis event. Data are mean±SEM. N=5 cells. **I.** Representative nucleokinesis events in a T cell migrating through a three-dimensional collagen matrix along a CCL19 chemokine gradient. Red arrows highlight full and partial nucleokinesis events. Time projections of the cellular and nuclear paths are shown in shades of green and blue, respectively. Time in hr:min:sec.

Full long-distance nuclear repositioning, also called nucleokinesis (Gundersen & Worman, 2013; Tsai & Gleeson, 2005), was rapid, with velocities of 2-9 micrometer per minute (Fig 1F), showed quick accelerations faster than the cell body (Fig 1H) and occurred over long intracellular distances of up to 45 micrometers (Fig 1G). To test whether nucleokinesis is a general feature of amoeboid immune cell migration, we imaged T cells, which have a smaller and less-branched cytoplasmic cell body, and still detected frequent and rapid nuclear repositioning from retracting into winning protrusions (Fig 1D-I, and Movie EV3). These findings identify nucleokinesis as a novel component of amoeboid immune cell migration.

### Nucleokinesis enables adaptive pathfinding in competing chemokine and pore size cues

As migrating cells encounter chemical (e.g., chemokine gradients) as well as mechanical (e.g., varying pores sizes) cues at the same time (Yamada & Sixt, 2019; Moreau *et al*, 2018; Kameritsch & Renkawitz, 2020), our observations suggested that cells may use nucleokinesis to flexibly adapt their path between locally competing guidance cues. To functionally test this hypothesis, we engineered reductionistic cellular path decision points, at which migrating cells encounter two path options with different strengths of chemotactic as well as pore size cues (Fig 2A-C). To test whether we can selectively guide cells to a specific path based on a stronger chemokine cue, we engineered one path to be closer to a chemokine source than the alternative path (Fig 2A). Indeed, when the two paths were composed of almost equally sized large pores at their path entrance, which are above the threshold of the nuclear pore size sensing mechanism, cells showed a strong preference to select the path that is closer to the chemokine source (Fig 2A and D, and Movie EV2). Yet, when we gradually narrowed the entrance pore of this particular path that is closer to the chemokine source, cells preferentially selected the alternative path that is more distant to the chemokine source but has a larger pore at the path entrance (Fig 2D-F, and Movie EV2). In contrast, in the absence of a chemokine source, cells always preferentially selected the longer paths with wider pores (Fig EV2A-C). Thus, these findings reveal that chemokine and pore size cues are competitive and identify thresholds at which chemokine cues overrule pore size cues and vice versa.

**Figure 2.**
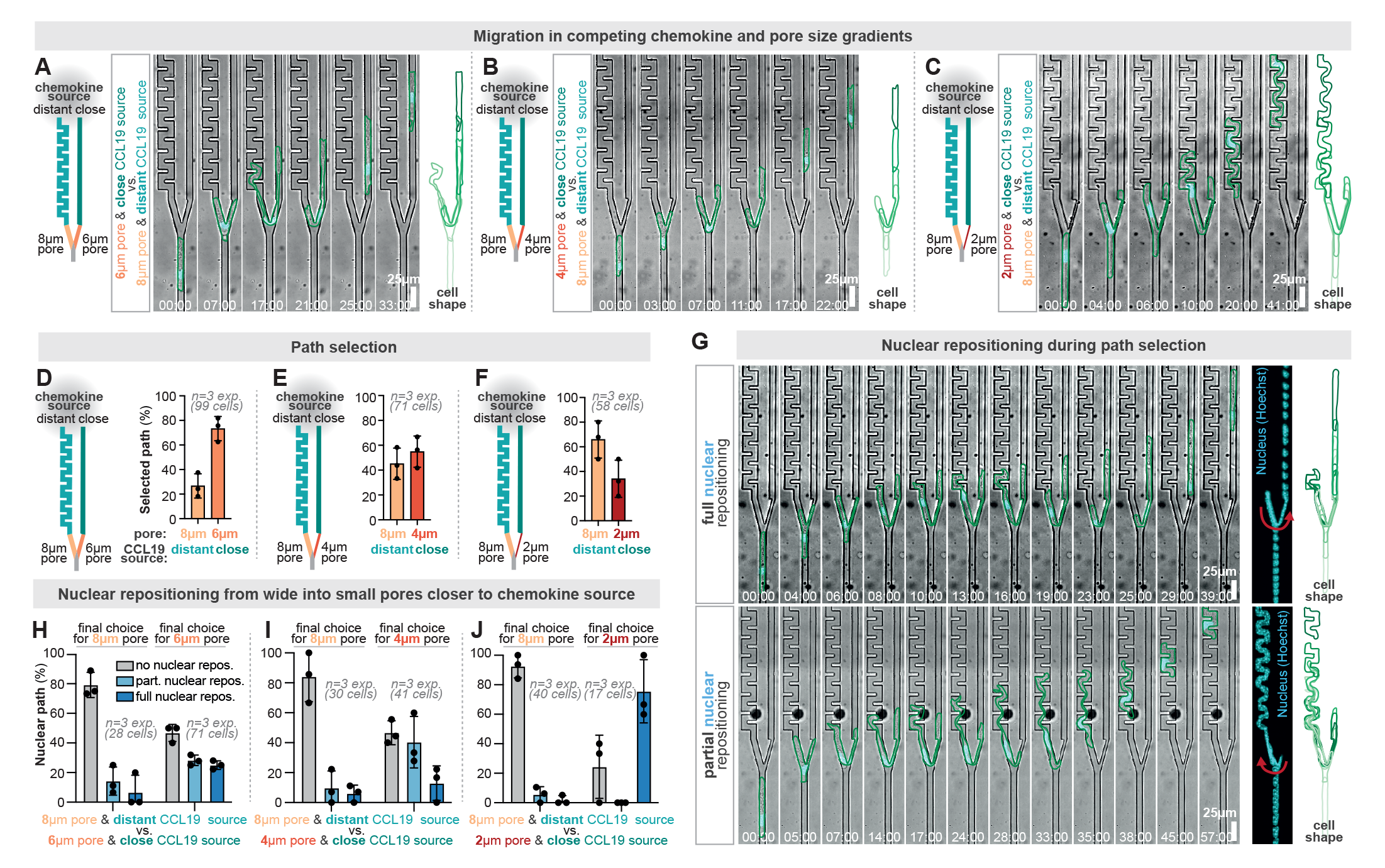
Nucleokinesis enables adaptive pathfinding in competing chemokine and pore size cues. **A.** Representative bone marrow-derived dendritic cell (DC) approaching a path junction with two alternative path options: the left path has an 8-micrometer wide pore, but is more distantly located to the CCL19 chemokine source. The right path has a 6-micrometer wide pore and is more closely located to the CCL19 chemokine source. The cell shape is outlined in green and the nucleus is visualized by Hoechst (cyan). Time in min:sec. **B.** As in A), but the right path that is closer to the chemokine source has a pore size of only 4- micrometer. **C.** As in A), but the right path that is closer to the chemokine source has a pore size of only 2- micrometer. **D.** Quantification of cellular path decisions in the microenvironments shown in A). See also Fig EV2A for controls without a chemokine source. Data are Mean±SD. N= 3 replicates (99 cells). **E.** Quantification of cellular path decisions in the microenvironments shown in B). See also Fig EV2B for controls without a chemokine source. Data are Mean±SD. N= 3 replicates (71 cells). **F.** Quantification of cellular path decisions in the microenvironments shown in C). See also Fig EV2C for controls without a chemokine source. Data are Mean±SD. N= 3 replicates (58 cells). **G.** Representative full and partial nucleokinesis events during cellular path decisions in the competing chemokine and pore sizes cues. The cell shape is outlined in green and the nucleus is visualized by Hoechst (cyan). Time in min:sec. **H.** Quantification of full and partial nucleokinesis events in the microenvironments shown in A), depending on whether the cell finally decides for the larger 8-micrometer or smaller 6- micrometer pore. Data are Mean±SD. N= 3 replicates (40 cells or 17 cells with the final decision either for 8- or 6-micrometer pore, respectively). **I.** Quantification of full and partial nucleokinesis events in the microenvironments shown in B), depending on whether the cell finally decides for the larger 8-micrometer or smaller 4- micrometer pore. Data are Mean±SD. N= 3 replicates (30 cells or 41 cells with the final decision either for 8- or 4-micrometer pore, respectively). **J.** Quantification of full and partial nucleokinesis events in the microenvironments shown in A), depending on whether the cell finally decides for the larger 8-micrometer or smaller 2- micrometer pore. Data are Mean±SD. N= 3 replicates (28 cells or 71 cells with the final decision either for 8- or 2-micrometer pore, respectively).

To test for the functional relevance of nucleokinesis, we imaged the spatiotemporal dynamics of nuclear behavior during these path decisions. While the nucleus was often immediately positioned in the future winning protrusion, we also observed frequent nucleokinesis events in which the nucleus was mispositioned far-distantly into the future ‘wrong’ path, necessitating long-distance nucleokinesis to the ‘winning’ protrusion (Fig 2G-J, and Movie EV2). Notably, nucleokinesis occurred particularly when the nucleus was initially positioned into the path bearing a larger pore entrance, but the cell selected the alternative path with a narrow pore entrance with a stronger chemotactic cue (Fig 2H-J). These findings show that cells can overcome the nuclear pore size sensing mechanism by chemotactic inputs and that nucleokinesis is required to adapt the nuclear path to the cellular path along an appearing dominating cue. Thus, nucleokinesis enables adaptive pathfinding during amoeboid cell navigation to flexibly navigate along locally competing mechanochemical guidance cues.

### Two-step amoeboid nucleokinesis by consecutive cell polarity switches

It is well established that cytoskeletal forces are able to intracellularly move and position the nucleus in diverse cell types such as neurons, glial cells, muscle cells, and fibroblasts (Calero-Cuenca *et al*, 2018; Cadot *et al*, 2015; Gundersen & Worman, 2013). Plotting the speed and distance of nucleokinesis in these cell types in comparison to the here measured parameters revealed an extraordinary efficiency of nucleokinesis in immune cells, being rapid as well as far distant (Fig EV1A). Notably, the intracellular behavior of the immune cell’s nucleus was reminiscent of the amoeboid’s cell body behavior, meaning that the nucleus moved rapidly, flexibly, and with continuous shape changes (e.g., Fig 1C and Movie EV1). As these parameters are hallmarks of amoeboid cell behavior, we propose to name nucleokinesis in amoeboid cells as ‘amoeboid nucleokinesis’.

To unravel the mechanistic basis of amoeboid nucleokinesis, we next imaged the spatiotemporal dynamics of the centrosome-to-nucleus axis, an important indicator of cell polarity and cytoskeletal forces acting on the nucleus during nuclear movement (Luxton & Gundersen, 2011). To uncouple nucleokinesis from effects caused by cellular squeezing, we established novel path junctions with equally large pore sizes but one blocked path, causing nucleokinesis events from the blocked to the open path (Fig EV2D-G, and Movie EV3). Using EB3-mCherry expressing DCs as a marker for the centrosome (also called microtubule-organizing center (MTOC)) and semi-automated imaging analysis (Fig EV2H), we observed two rapid polarity switches in the nucleus-MTOC axis configuration during amoeboid nucleokinesis (Movie EV3): while DCs migrated with a typical amoeboid nucleus-forward configuration (Renkawitz *et al*, 2019) before nucleokinesis (Fig 3A-C), the re-positioning of the nucleus from the ‘losing’ into the ‘winning’ protrusion coincided with a switch in the nucleus-MTOC axis configuration as the cells moved with the MTOC in front of the nucleus during the first phase of nucleokinesis (Fig 3A-C). When the DCs continued their path of migration upon a successful nucleokinesis event, the nucleus passed again the MTOC to restore the initial nucleus-MTOC axis configuration (Fig 3A-C), demonstrating that the nucleus is not only passively dragged by retracting protrusions but actively repositions to its nucleus-forward configuration.

**Figure 3.**
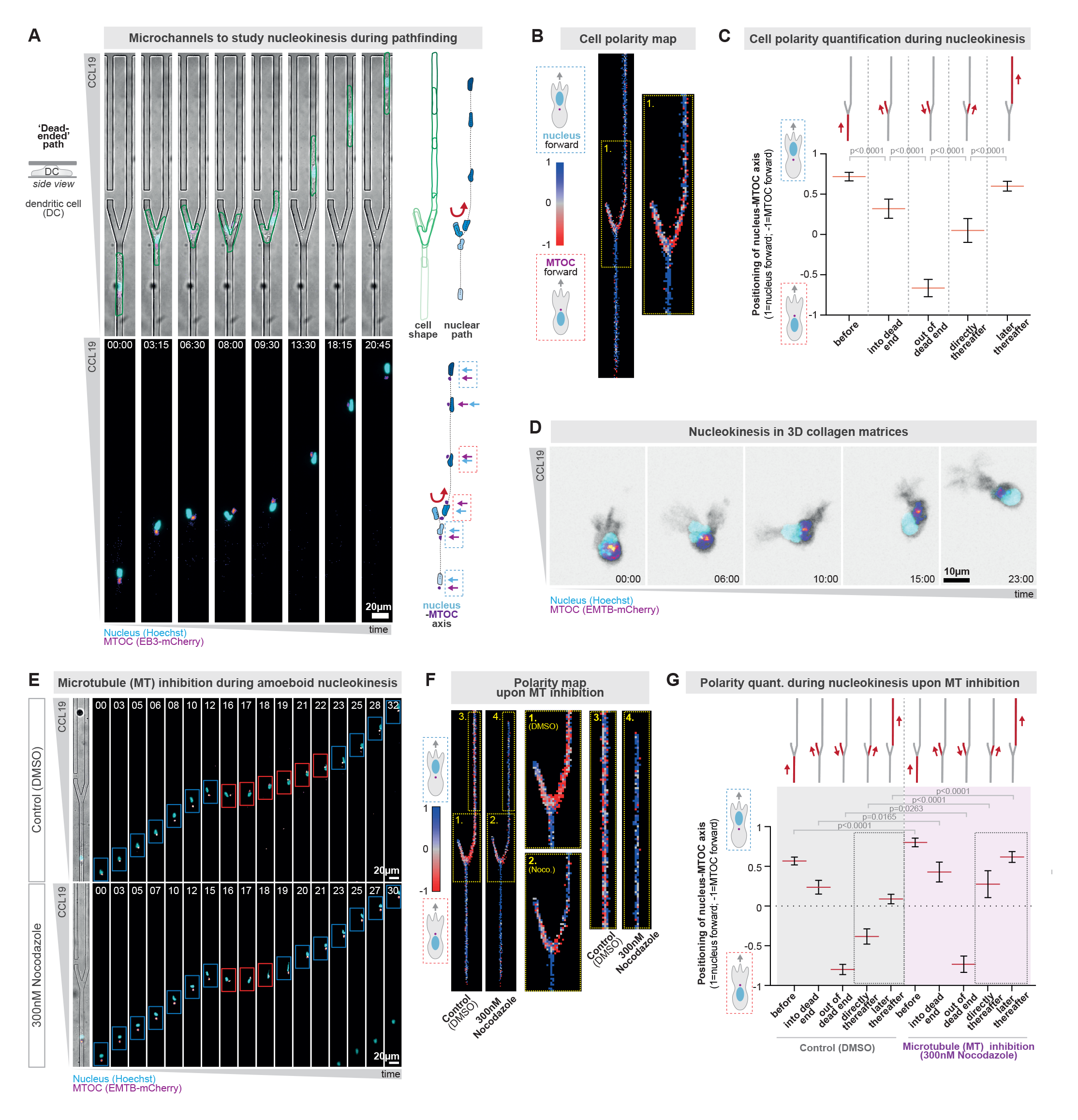
Two-step amoeboid nucleokinesis by consecutive cell polarity switches. **A.** Representative Hoxb8-derived dendritic cell (DC) approaching a path decision with equal pore sizes but one blocked path, frequently causing nucleokinesis from the blocked to the open path. The DC stably encodes the microtubule plus-end marker EB3-mCherry, which also visualizes the microtubule-organizing center (MTOC; in pink). The nucleus is visualized by Hoechst (cyan) and the cell shape is outlined in green. Projections of cellular (green), MTOC (pink) and nuclear (blue) paths are shown on the right. The configuration of the nucleus-MTOC axis is highlighted by dashed boxes (blue=nucleus forward; red=MTOC forward). See also Fig EV2D-G for a detailed characterization of the assay. Time in min:sec. **B.** Heatmap of the nucleus-MTOC axis configuration during amoeboid nucleokinesis. The frontward positioning of the nucleus is depicted in blue and the frontward positioning of the MTOC is depicted in red. The yellow-dotted region is shown enlarged to depict the cellular behavior during initial nucleokinesis. See also Fig EV2H for a more detailed description of the imaging quantification. N=6 replicates, 48 cells. **C.** Quantification of the nucleus-MTOC axis configuration before, during, directly after, and later after amoeboid nucleokinesis (1=all cells position the nucleus in front of the MTOC; −1= all cells position the MTOC in front of the nucleus). Data are mean±95CI, Mann-Whitney test, N=6 replicates, 48 cells, and 660 (before), 250 (into a dead end), 191 (out of a dead end), 179 (directly thereafter), and 678 (later thereafter) image frames. **D.** Representative HoxB8-derived DCs stably encoding EMTB-mCherry (visualizes the MTOC; fire-color coded) and transiently stained with Hoechst (visualizes the nucleus; cyan), migrating in a three-dimensional collagen network along a CCL19 chemokine gradient. The cell shape is shown in black (via the high intensity of the EMTB-mCherry channel). Note the nucleokinesis event from the left protrusion into the right protrusion. Time in min:sec. **E.** Representative HoxB8-derived DCs in the presence of the microtubule inhibitor nocodazole (300nM) or control (DMSO). The DCs stably encode EMTB-mCherry (to visualize the MTOC; fire-color coded) and is transiently stained with Hoechst (to visualize the nucleus; cyan). The configuration of the nucleus-MTOC axis is highlighted by boxes (blue=nucleus forward; red=MTOC forward). Time in min. **F.** Heatmap of the nucleus-MTOC axis configuration during amoeboid nucleokinesis in the presence of the microtubule inhibitor nocodazole (300nM) or control (DMSO). The yellow-dotted regions 1 (DMSO) and 2 (Nocodazole) are enlarged to depict the cellular behavior during the initial nucleokinesis event, and the yellow-dotted regions 3 (DMSO) and 4 (Nocodazole) are enlarged to depict the cellular behavior during the later nucleokinesis events to reposition the nucleus to the cellular front. N=3 replicates, 96 (DMSO) and 36 (Nocodazole) cells. **G.** Quantification of the nucleus-MTOC axis configuration before, during, directly after, and later after amoeboid nucleokinesis in the presence of the microtubule inhibitor nocodazole (300nM) or control (DMSO) (1=all cells position the nucleus in front of the MTOC; −1= all cells position the MTOC in front of the nucleus). Data are mean±95CI, Mann-Whitney test, N=3 replicates, 96 (DMSO) and 36 (Nocodazole) cells, and 1096 (before; DMSO), 463 (before; nocodazole), 493 (into a dead end, DMSO), 203 (into a dead end, nocodazole), 338 (out of a dead end, DMSO), 172 (out of a dead end, nocodazole), 360 (directly thereafter, DMSO), 127 (directly thereafter, nocodazole), 1062 (later thereafter, DMSO), and 508 (later thereafter, nocodazole) image frames.

To corroborate these findings, we used the microtubule markers EMTB-mCherry and Spy-tubulin as additional MTOC markers (Fig EV3A and B), as well as manual mapping of the MTOC-to-nucleus axis during nucleokinesis (Fig EV3C-E, and Fig EV4A-E), confirming both observed rapid polarity switches during amoeboid nucleokinesis. Similarly, when we investigated the MTOC-to-nucleus axis during DC pathfinding in deformable collagen networks, we observed preferential MTOC-forward configuration during the first step of nucleokinesis, followed by the restoration of the nucleus-forward configuration (Fig 3D). Together, these findings identify that amoeboid nucleokinesis is characterized by two rapid polarity switches in the MTOC-to-nucleus axis.

### Myosin-based pushing drives amoeboid nucleokinesis to ensure adaptive pathfinding

The positioning of the MTOC in front of the nucleus during the initial phase of nuclear repositioning raised the possibility that microtubules emanating from the MTOC may couple to the nucleus, and thereby exert forces on the nucleus to pull it from the ‘losing’ protrusion back to the cellular path. Yet surprisingly, when we depolymerized microtubules (Fig EV5A and B), we did not observe any slowdown of amoeboid nucleokinesis (Fig 3E, Fig EV5C and D, and Movie EV4). In contrary, nocodazole-treated cells showed an accelerated speed of nucleokinesis (Fig EV5C and D) as well as an accelerated nuclear repositioning frontward to the MTOC (Fig 3F and G). As microtubule depolymerization is well known to cause increased cellular contractility by releasing microtubule-bound actomyosin regulators like the RhoA guanine nucleotide exchange factor GEF-H1 (Kopf *et al*, 2020; Bouchet & Akhmanova, 2017; Krendel *et al*, 2002), this finding raised the possibility that actomyosin contractility could be a driving force of amoeboid nucleokinesis. This notion was supported when we quantified the distance as well as the change of distance between the nucleus and the MTOC (Fig EV4D and E), showing that the nucleus is even more distantly frontward positioned upon microtubule inhibition (Fig EV5E and F), as if the nucleus would be pushed to a frontward localization while the localization of the MTOC remains rather stable along the cell axis. Further, imaging of myosin-IIA localization during amoeboid nucleokinesis using MYH9-GFP encoding dendritic cells revealed an enriched myosin-IIA localization in retracting protrusions closely behind the nucleus (Fig 4A and B, and Movie EV5).

**Figure 4.**
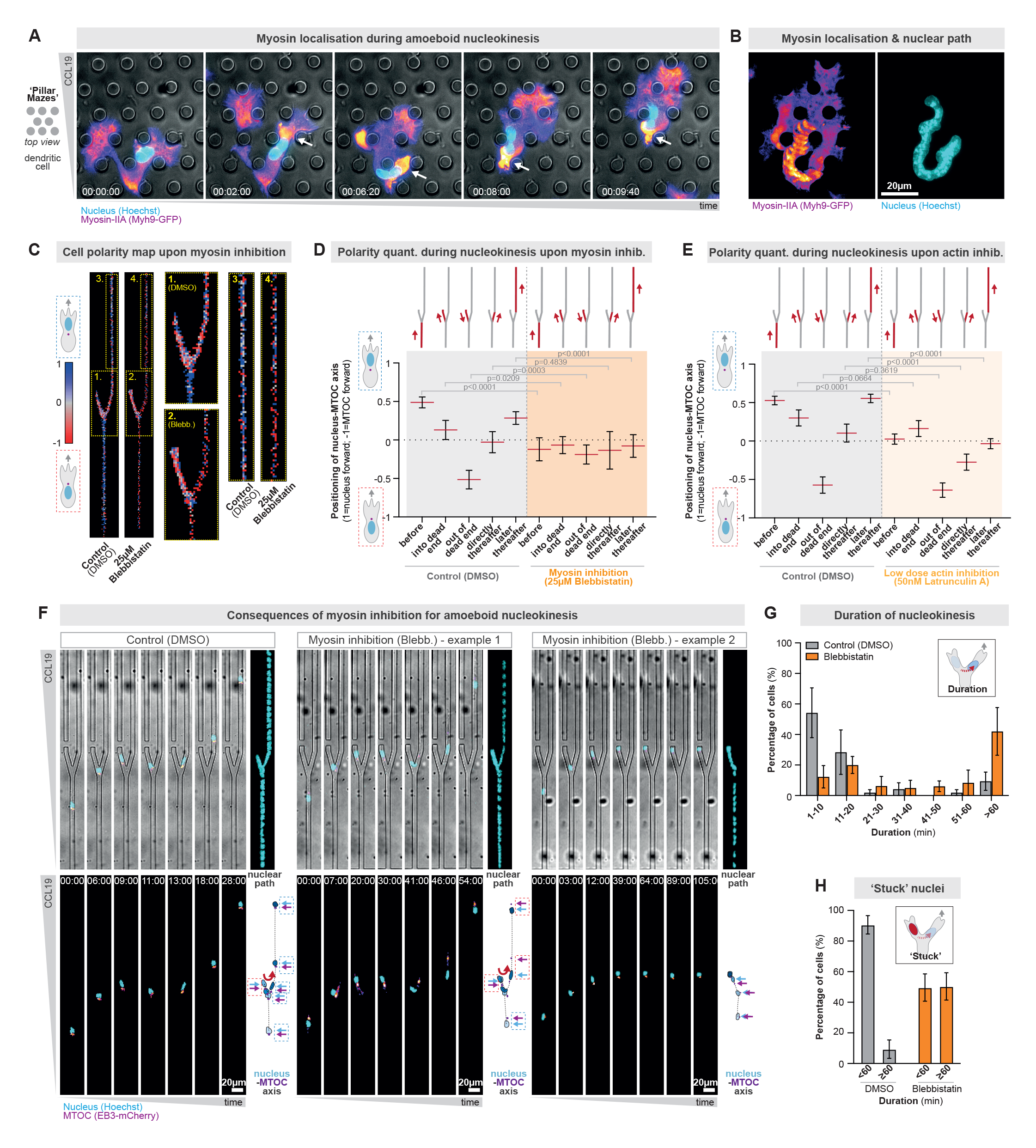
Myosin-based pushing drives amoeboid nucleokinesis to ensure adaptive pathfinding. **A.** Representative HoxB8-derived Myh9-GFP expressing DC migrating through a porous maze-like microenvironment composed of pillars interconnecting two surfaces below and above the migrating cell. Myh9-GFP localization is fire color-coded and the nuclear localization is shown in cyan (transient Hoechst label). Time in hr:min:sec. **B.** Time projection of Myh9-GFP and nuclear localization of the cell shown in A). **C.** Heatmap of the nucleus-MTOC axis configuration during amoeboid nucleokinesis in the presence of the myosin-II inhibitor para-nitroblebbistatin (25μM) or control (DMSO). The yellow-dotted regions 1 (DMSO) and 2 (para-nitroblebbistatin) are enlarged to depict the cellular behavior during the initial nucleokinesis event, and the yellow-dotted regions 3 (DMSO) and 4 (para-nitroblebbistatin) are enlarged to depict the cellular behavior during the later nucleokinesis events to reposition the nucleus to the cellular front N=4 replicates, 37 (DMSO) and 13 (para-nitroblebbistatin) cells. **D.** Quantification of the nucleus-MTOC axis configuration before, during, directly after, and later after amoeboid nucleokinesis in the presence of the myosin-II inhibitor para-nitroblebbistatin (25μM) or control (DMSO) (1=all cells position the nucleus in front of the MTOC; −1= all cells position the MTOC in front of the nucleus). Data are mean±95CI, Mann-Whitney test, N=4 replicates, 37 (DMSO) and 13 (para-nitroblebbistatin) cells, and 582 (before; DMSO), 173 (before; para-nitroblebbistatin), 246 (into a dead end, DMSO), 315 (into a dead end, para-nitroblebbistatin), 198 (out of a dead end, DMSO), 244 (out of a dead end, para-nitroblebbistatin), 208 (directly thereafter, DMSO), 67 (directly thereafter, para-nitroblebbistatin), 523 (later thereafter, DMSO), and 180 (later thereafter, para-nitroblebbistatin) image frames. **E.** Quantification of the nucleus-MTOC axis configuration before, during, directly after, and later after amoeboid nucleokinesis in the presence of the actin inhibitor LatrunculinA (50nM) or control (DMSO) (1=all cells position the nucleus in front of the MTOC; −1= all cells position the MTOC in front of the nucleus). Data are mean±95CI, Mann-Whitney test, N=3 replicates, 78 (DMSO) and 13 (LatrunculinA) cells, and 900 (before; DMSO), 866 (before; LatrunculinA), 323 (into a dead end, DMSO), 351 (into a dead end, LatrunculinA), 234 (out of a dead end, DMSO), 265 (out of a dead end, LatrunculinA), 281 (directly thereafter, DMSO), 328 (directly thereafter, LatrunculinA), 902 (later thereafter, DMSO), and 899 (later thereafter, LatrunculinA) image frames. **F.** Representative HoxB8-derived DCs in the presence of the myosin-II inhibitor para-nitroblebbistatin (25μM) or control (DMSO). The DCs stably encode EB3-mCherry (to visualize the MTOC; fire color-coded) and are transiently stained with Hoechst (to visualize the nucleus; cyan). The left panel shows a representative control cell, the middle panel a representative myosin-inhibited cell with delayed nuclear repositioning to the front of the cell, and the right panel a representative myosin-inhibited cell that entirely fails to reposition to the productive open path. The configuration of the nucleus-MTOC axis is highlighted by dashed boxes (blue=nucleus forward; red=MTOC forward). Time in min:sec. **G.** Quantification of the duration of nucleokinesis in the presence of the myosin-II inhibitor para-nitroblebbistatin (25μM) or control (DMSO). Data are mean±SEM. N=3 replicates, 50 (DMSO) and 38 (para-nitroblebbistatin) cells. **H.** Quantification of the percentage of cells that have a nucleus stuck in the blocked path for at least 60 minutes (myosin-II inhibitor para-nitroblebbistatin (25μM) versus control (DMSO)). Data are mean±SEM. N=3 replicates, 50 (DMSO) and 38 (para-nitroblebbistatin) cells.

To test whether myosin-IIA indeed provides forces to reposition the nucleus during amoeboid nucleokinesis, we next exposed motile DCs to the myosin-II inhibitor para-nitroblebbistatin (Képiró *et al*, 2014) (Fig. EV6A and B). Cells with inhibited myosin failed to efficiently switch cell polarities during nucleokinesis, showing a random nucleus-to-MTOC axis configuration before, during, and after nucleokinesis (Fig 4C and D, Fig EV6C-G, and Movie EV6). To corroborate these results, we analyzed myosin-IIA knockout DCs and also observed delayed reconfiguration of the nucleus-to-MTOC axis during amoeboid nucleokinesis (Fig EV6H). Given the concerted action of myosin together with the actin cytoskeleton, we next inhibited the actin cytoskeleton with low-doses of latrunculin, which still enabled migration (Fig EV7A and B) but reduced the rate of cell polarity switching during nuclear repositioning (Fig 4E and EV7A-I). Thus, actomyosin contractility is required for efficient cell polarity switching during amoeboid nucleokinesis. To test for the functional consequences of impaired amoeboid nucleokinesis, we next measured the speed of nucleokinesis and cellular path adaptions when myosin is non-functional. While the cellular speed before nucleokinesis was even mildly increased (Fig EV6E), repositioning of the nucleus from the ‘dead’-end path to the selected cellular path was strongly slowed down (Fig 4G, Fig EV6D and E, and Fig EV7E-G), causing a longer duration of nucleokinesis (Fig 4F and G) and even entire failure to adapt the nuclear to the cellular path, resulting in cells stuck in the wrong path (Fig 4F and H, and Movie EV6). Together, these findings identify actomyosin forces as a major driver of amoeboid nucleokinesis.

### Adaptive pathfinding by nucleokinesis in the *Dictyostelium discoideum* amoebae

To explore whether adaptive pathfinding by amoeboid nucleokinesis is a general feature of amoeboid cell migration, we investigated motile single cells of the amoebae *Dictyostelium discoideum*. Comparable to immune cells, *Dictyostelium* cells typically explore their surrounding microenvironment with at least two protrusions (Andrew & Insall, 2007), until they select a path, followed by repeated cycles of protrusion formation, path exploration, and path selection (Fig 5A, and Movie EV7). Using a nuclear marker during *Dictyostelium* pathfinding in an environmental maze along a chemotactic folate gradient, we discovered frequent nucleokinesis events from future ‘losing’ into ‘winning’ protrusions (Fig 5A and E), with rapid speeds faster than the cell front (Fig 5B) in the range of 8 to 25 micrometer per minute (Fig 5C). Given the smaller cell body of *Dictyostelium* cells in comparison to DCs, the nucleokinesis distances of 10 micrometer represented far-distant intracellular nuclear re-positioning throughout the cell body (Fig 5D). Thus, the basic parameters of nucleokinesis in *Dictyostelium* cells are highly comparable to nucleokinesis in immune cells (Fig 5C-E, Fig EV1A-C, and Movie EV7). Next, to characterize the configuration of the nucleus-MTOC axis, we employed *Dictyostelium* cells encoding mRFP-histone as a marker for the nucleus and GFP-α-tubulin as a marker for the MTOC (Fig 5F). Before a nucleokinesis event, *Dictyostelium* cells migrated in the typical amoeboid nucleus-forward configuration (Ishikawa-Ankerhold *et al*, 2022), but then switched to an MTOC-forward configuration during nucleokinesis, followed by the restoration of the nucleus-forward configuration (Fig 5F-H, and Movie EV8). Notably, as in amoeboid immune cells, this two-step polarity switch is dependent on myosin-II contractility, as the positioning of the nucleus to the front of the cell axis was strongly delayed in the presence of the myosin-II inhibitor para-nitroblebbistatin (Fig 5I, Fig EVA and B, and Movie EV9). To corroborate these results, we analyzed *Dictyostelium* cells bearing a well-established myosin-II null mutation (myosin-II deficient (mhcA-null) strain HS2205) (Manstein *et al*, 1989; Bindl *et al*, 2020), as well as genetically encoded markers for the nucleus and the MTOC (mRFP-histone and GFP-α-tubulin), and also observed delayed nuclear relocalization frontward to the MTOC during nucleokinesis (Fig 5J and K, and Movie EV9). Thus, despite the phylogenetic distance between mammalian immune cells and the amoebae *Dictyostelium discoideum*, the mechanistic principles of amoeboid nucleokinesis during cellular navigation are conserved.

**Figure 5.**
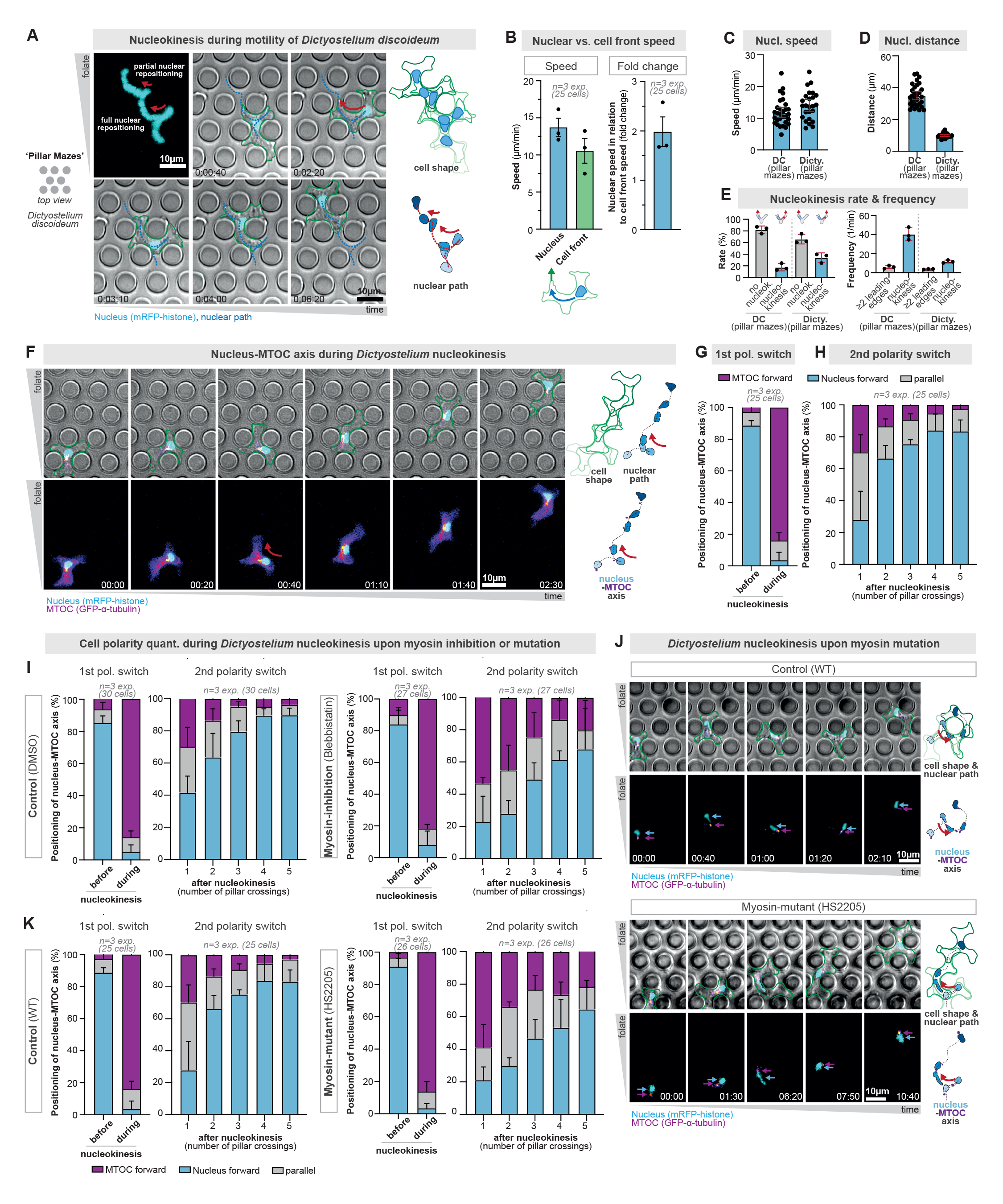
Adaptive pathfinding by nucleokinesis in *Dictyostelium discoideum* amoebae. **A.** Representative *Dictyostelium discoideum* cell migrating through a porous maze-like microenvironment composed of pillars interconnecting two surfaces below and above the migrating cell. The *Dictyostelium* cell stably expresses mRFP-histone (to visualize the nucleus; cyan) and the cell shape is outlined in green. Time projections of the cellular and nuclear paths are shown in shades of green and blue, respectively. Time in hr:min:sec. **B.** Quantification of the speed of nuclear and cell front movement during full nucleokinesis in *Dictyostelium* cells. Independent speeds of nucleus and cell front are shown on the left. Nuclear speed normalized to the cell front speed (fold change) is shown on the right. Data are mean±SD. N=3 replicates, 25 cells. **C.** Quantification of the speed of nuclear movement during nucleokinesis in DCs and *Dictyostelium* cells migrating in pillar mazes (DCs in pillars mazes with 9 micrometer pores sizes, and *Dictyostelium* cells in pillar mazes with 5 micrometer pore sizes). Data are median±95CI. N=3 replicates, 25 (*Dictyostelium*) and 30 (DCs) cells. **D.** Quantification of the intracellular distance of nuclear movement during nucleokinesis in DCs and *Dictyostelium* cells migrating in pillar mazes. Data are median±95CI. N=3 replicates, 25 (*Dictyostelium*) and 30 (DCs) cells. **E.** Quantification of the rate (left) and frequency (right) of nucleokinesis events in amoeboid migrating *Dictyostelium* cells and DCs. Rate: when the migrating cell has a least two major protrusions, quantification of whether the nucleus locates immediately into the dominant protrusion or first locates into the future retracting protrusion, requiring nucleokinesis into the future dominant protrusion. Frequency: showing the frequency of at least two leading edges and nucleokinesis events per minute. Data are mean±SD. N=3 replicates, 25 (*Dictyostelium*) and 30 (DCs) cells. **F.** Representative *Dictyostelium* cell migrating in a pillar maze. The *Dictyostelium* cell stably encodes GFP-α-tubulin, which also visualizes the microtubule-organizing center (MTOC; in pink) and mRFP-histone, which visualizes the nucleus (in cyan). The cell shape is outlined in green. Projections of cellular (green), MTOC (pink) and nuclear (blue) paths are shown on the right. Red arrows highlight full and partial nucleokinesis events. Time in min:sec. **G.** Quantification of the nucleus-MTOC axis configuration before and during nucleokinesis. Data are mean±SD. N=3 replicates, 25 cells. **H.** Quantification of the nucleus-MTOC axis configuration after nucleokinesis. The x-axis indicates the number of pillar crossings after nucleokinesis. Data are mean±SD. N=3 replicates, 25 cells. **I.** Quantification of the nucleus-MTOC axis configuration before, during, and after nucleokinesis upon myosin inhibition. Data are mean±SD. N=3 replicates, 30 (DMSO) and 27 (para-nitroblebbistatin) cells. **J.** Representative *Dictyostelium* cells (WT: top, HS2205 myosin mutant: bottom) migrating in a pillar maze. The *Dictyostelium* cells stably encode GFP-α-tubulin, which also visualizes the microtubule-organizing center (MTOC; in pink) and mRFP-histone, which visualizes the nucleus (in cyan). The cell shape is outlined in green. Projections of cellular (green), MTOC (pink) and nuclear (blue) paths are shown on the right. Red arrows highlight full and partial nucleokinesis events. Time in min:sec. **K.** Quantification of the nucleus-MTOC axis configuration before, during, and after nucleokinesis upon myosin mutation. Data are mean±SD. N=3 replicates, 25 (WT) and 26 (HS2205; myosin mutant) cells.

Together, these findings identify amoeboid nucleokinesis as a conserved process in rapid amoeboid migrating cells, comprising a two-step cell polarity switch driven by myosin-II forces, sliding the nucleus from a ‘losing’ to the ‘winning’ protrusion to ensure adaptive pathfinding in complex microenvironments.

## Discussion

Cellular organelles are often actively positioned to defined subcellular locations within the cytoplasm (Bornens, 2008). This active positioning includes membrane-surrounded organelles like the nucleus and mitochondria, but also membraneless-organelles like the microtubule-organizing center (MTOC). The active positioning of the genome-carrying nucleus is particularly challenging (Gundersen & Worman, 2013), as it is typically the largest and stiffest organelle (Kalukula *et al*, 2022). Cells solved this challenge by exerting forces from the cytoskeleton onto the nucleus, leading to its intracellular movement and positioning, a process termed nucleokinesis. For instance, cells in the developing vertebrate neuroepithelium employ forces that are mainly generated by the actomyosin cytoskeleton to move their nuclei basally and apically during the progression of the cell cycle (Norden *et al*, 2009), whereas the movements of female and male pronuclei during fertilization are mainly driven by forces from the microtubule cytoskeleton (Reinsch & Gonczy, 1998).

An additional complexity for nuclear positioning arises when cells are not stationary but motile (Calero-Cuenca *et al*, 2018), adding the challenge of coordinating nuclear positioning with the cellular advance along the migratory path. Key findings showed that motile fibroblasts reposition their nucleus to the cellular rear to start their movement into cell-free tissue wounds (Zhu *et al*, 2018; Luxton *et al*, 2010). Mechanistically, this rearward nuclear positioning is driven by an actin cortex composed of actin cables that are coupled to the nucleus and move against the direction of cellular movement (Luxton *et al*, 2010). Fibroblasts are mesenchymally migrating cells that adhere to their extracellular environments and migrate with velocities in the range of tens of micrometers per hour. Thus, the principles and mechanisms of nuclear positioning in stationary and slowly moving cells are increasingly well understood. However, how extremely fast migrating cells coordinate the intracellular positioning of the nucleus with their migration path remained entirely unknown. The high velocities of fast migrating cells, which are typically around ten micrometers per minute, even raised the question of whether nuclear movement can be at all faster than the cellular speed, which would be required for active nuclear repositioning. On top of this speed challenge, rapidly migrating cells also provide a morphological challenge: the fastest migrating cells are typically amoeboid cells, including many immune cells, which have complex, ramified cell shapes that dynamically and constantly change during navigation along the migration path, raising the question whether and how nuclear positioning is coordinated with these highly dynamic changes in cell shape.

Here, we investigated major cellular models of fast amoeboid migrating cells, including immune cells and the single-cell amoeba *Dictyostelium discoideum*, and discovered that motile amoeboid cells actively move and position their nuclei during pathfinding. Nuclear movement in amoeboid cells is extraordinarily efficient and frequent, functioning in a very rapid manner over long intracellular distances. Given the similarity of this nuclear movement to the movement properties of entire amoeboid cells, we here name this newly described mode of nuclear movement as ‘amoeboid nucleokinesis’. Amoeboid nucleokinesis functions independently of microtubules but requires forces from the actomyosin cytoskeleton. The surprisingly fast temporal scale of nuclear repositioning and of the two switches in the configuration of nucleus-MTOC axis, raise the intriguing model that amoeboid nucleokinesis might be solely driven by myosin pushing forces from behind the nucleus, moving the nucleus forward without any requirement of anchorage to the cytoskeleton. Functionally, we discover that amoeboid nucleokinesis is required for amoeboid cell migration to enable navigation in complex mechanochemical microenvironments. Specifically, amoeboid nucleokinesis allows cells to flexibly adapt their path to an appearing dominant guidance cue. Considering that virtually all motile cells simultaneously encounter a variety of chemical and mechanical signals in their immediate local microenvironment, these findings suggest that nucleokinesis plays a universally crucial role in the navigation of migrating cells.

## Materials and Methods

### Cell culture

#### Mammalian cell culture

All cells were kept at 37°C in a humidified incubator with 5 % CO2. DCs were differentiated either from the bone marrow of male C57B16/J wildtype mice (aged 8-12 weeks), male or female MyoIIA-Flox*CD11c-Cre mice (aged 8-11 weeks), or from Hoxb8 precursor lines (EB3-mCherry, EMTB-mCherry, and Myh9-GFP). Cells were differentiated in R10 medium (RPMI 1640 supplemented with 10 % fetal calf serum (FCS), 2 mM L-glutamine, 100 U/ml penicillin, 100 mg/ml streptomycin and 0.1 mM 2- mercaptoethanol; all Gibco) supplemented with 10 % granulocyte-macrophage colony-stimulating factor (GM-CSF) hybridoma supernatant. Fresh medium was added on differentiation days 3 and 6. To induce maturation, either fresh or thawed DCs (differentiation day 8) were stimulated for 24 h with 200 ng/ml lipopolysaccharide (LPS; *E. coli* O26:B6, Sigma-Aldrich) and used for experiments on day 9. T cells were isolated from the spleen of male or female C57BL/6J mice (aged 6-12 weeks) using the EasySep mouse naive T cell isolation Kit (Stemcell). Cells were seeded onto cell culture plates coated with 1 µg/ml CD3 antibody and 1 µg/ml CD28 antibody and either used for experiments between differentiation days 3 and 6 or frozen at day 6 and thawed for experiments.

#### Dictyostelium discoideum

Cells of the *D. discoideum* strain AX2-214 (here designated as wild type), and the myosin-II deficient (mhcA-null) strain HS2205 derived from it (Manstein *et al*, 1989) were used. Nuclei and microtubules in both strains were visualized by expression of GFP-α-tubulin (tubA1; DDB0191380|DDB_G0287689) and mRFP-histone (H2Bv3; DDB0231622|DDB_G0286509) (Ishikawa-Ankerhold *et al*, 2022; Bindl *et al*, 2020). Cells were cultured in polystyrene Petri dishes containing HL5 medium (Formedium, Hunstanton, Norfolk, UK) supplemented with 10 µg/ml of blasticidin S (Gibco, Fisher Scientific GmbH, Schwerte, Germany), and 20 µg/ml of geneticin (Sigma-Aldrich, Sigma-Aldrich Chemie GmbH, Taufkirchen, Germany) or 33 µg/ml hygromycin B (EMD Millipore Corp., Billerica, MA, USA) at 22°C. While the myosin-II deficient mutant has been reported to have cytokinesis defects causing multi-nucleation, our adhesive cell culture conditions mostly resulted in single-nucleated cells and for analysis we only included those single-nucleated cells.

### Mice

Wild-type animals were housed in the Core Facility Animal Models at the Biomedical Centre (Ludwig-Maximilians-Universität) and animal procedures and experiments were in accordance with the ministry of animal welfare of the region of Oberbayern and with the German law of animal welfare.

### Flow cytometry analysis

DCs were routinely checked for surface marker expression using antibodies for CD11c (17- 0114-82, Invitrogen) and MHCII (48-5321-82, Invitrogen). After Fc receptor blockage using an antibody for CD16/32 (14-0161-85, Invitrogen), stainings were performed in FACS buffer (1 % BSA, 2 mM EDTA in PBS). Cells were analyzed using a Cytoflex S flow cytometer (Beckmann-Coulter).

### Micro-fabricated devices

Micro-fabricated devices were generated as described previously (Kroll *et al*, 2022; Renkawitz *et al*, 2018). In brief, wafers produced by photolithography or epoxy replicates thereof were used as templates for micro-structures with defined lengths, widths, and heights. Micro-channels had a width of 8 µm. For analysis of adaptive pathfinding in competing chemokine and pore-size cues, pores were 2, 4 or 6 µm wide. Pore sizes in pillar forests were 5 and 9 µm for *Dictyostelium discoideum* and DC migration, respectively. The height of the micro-structures ranged from 4-5 µm to ensure cell confinement from all sides.

Polydimethylsiloxane (PDMS; 10:1 mixture of Sylgard 184, Dow) was cast on the template micro-structures. PDMS was mixed in a Thinky mixer, and air bubbles were removed in a desiccator. After curing at 80°C overnight, PDMS was removed from the template and cut into single devices. Holes were punched on each side of the micro-structures to enable cell and chemokine loading. Subsequently, PDMS pieces were bonded to cleaned coverslips using a plasma cleaner. PDMS devices were placed at 120°C for 10 min and at 80°C overnight to ensure permanent bonding.

### Live-cell migration assays

To visualize the nucleus for live-cell migration assays, cells were incubated with 1 drop of NucBlue (Invitrogen) in 1ml of cell/media mixture for at least 30 min. SPY555-tubulin (Spirochrome) was used according to the manufacturer’s protocol to visualize the MTOC. For pharmacological inhibition experiments, cells were treated with final concentrations of 50 nM Latrunculin A (Sigma-Aldrich), 25 µM para-nitroblebbistatin (Motorpharma; dissolved in DMSO), or 300 nM Nocodazole (Sigma-Aldrich). Control samples were treated with the respective DMSO dilution.

#### Microchannel migration assays

PDMS devices were flushed with phenol-free R10 medium supplemented with 50 µM L-ascorbic acid (Sigma-Aldrich) and pharmacological inhibitors if applicable for the experimental setup. Devices were incubated at 37°C, 5 % CO2 for at least 1 hour before the experiment. Subsequently, 0.625 µg/ml CCL19 (DCs) or 100 µM folate (*Dictyostelium*) were loaded into the chemokine loading hole to establish a gradient. Finally, 0.1-1×10^5^ cells were added into the opposite loading hole.

#### Collagen migration assays

Collagen migration assays were performed as described previously (Kroll *et al*, 2022). Briefly, for DC or T cell migration in collagen, 1x minimum essential medium (MEM, Gibco), 0.4% sodium biocarbonate (Sigma-Aldrich) and Nutragen bovine collagen (Advanced BioMatrix) were mixed with 3-5×10^5^ cells in R10 medium at a 2:1 ratio, resulting in a final collagen density of 3.3 mg/ml. The collagen-cell solution was added into custom-made migration chambers (appr. 17 mm in width and 1 mm in height). After 75 min polymerization at 37°C, 5 % CO2, gels were overlaid with 80 µl CCL19 (0.625 µg/ml). For pharmacological inhibition experiments, inhibitors were added to the chemokine solution as well as the collagen-cell solution.

### Imaging

Live-cell imaging of dendritic cells and T cells was performed at 37°C and with 5 % CO2 in a humidified chamber. Live-cell imaging of *Dictyostelium discoideum* was conducted at room temperature (22°C). Cell migration was recorded using inverted DMi8 microscopes (Leica) with HC PL FLUOTAR 4x/0.5 PH0 air, HC PL APO 20x/0.80 PH2 air, or HC PL APO 40x/0.95 CORR air objectives. Additionally, the microscope was equipped with an LED5 (Leica) or pE-4000 (CoolLED) light source, an incubation chamber, a heated stage, and a CO2 mixer (Pecon).

### Image analysis

Fiji/ImageJ (Schindelin *et al*, 2012) and Imaris (Bitplane) were used for image processing. Generally, only single, non-interacting cells were used for quantification to exclude the influence of neighboring cells on cell path, speed, pore size decision or the nucleus-MTOC axis.

The overall speed of cells in collagen matrices was analyzed using a custom-made cell tracking tool for ImageJ (Kiermaier *et al*, 2016). In brief, image sequences were background corrected, and particle filtering was used to exclude objects larger or smaller than cells. Each image of the sequence was matched with the optimal overlap in its lateral displacement to the previous frame. Finally, migration velocity was calculated from the y-displacement and the time between two consecutive frames. Speed and distance of nucleus and MTOC in single migrating DCs, T cells, and *Dictyostelium* cells in collagen matrices as well as in pillar forests were analyzed using the manual tracking plugin in Fiji (v2.1.1). Orientation of nucleus and MTOC, as well as nuclear repositioning (whole nuclear body located in losing protrusion followed by full repositioning into winning protrusion), was quantified manually in Fiji.

For the analysis of cells migrating in 1x dead end-channels, the nuclei were tracked using the tracking function of Imaris v9.7.2 with the following settings: object diameter 10 µm, manually adjusted quality threshold, autoregressive motion tracking algorithm (max. distance 20 µm, gap size=1). MTOC signals were either tracked manually in Imaris (SPY555-tubulin and EMTB-mCherry) or segmented in ilastik 1.4.0 (Berg *et al*, 2019) (EB3-mCherry) and then tracked using the tracking function of Imaris v9.7.2 with the following settings: object diameter 3 µm, manually adjusted quality threshold, autoregressive motion tracking algorithm (max. distance 25 µm, gap size=3). Subsequently, all automated tracking was manually evaluated for errors. Position data for nuclei and MTOCs were exported and analyzed by a custom-made Matlab script. To accurately associate MTOC tracks with corresponding nuclei, track pairing is achieved by minimizing the convex hull volume between their points. For each nuclear track, the motion direction is calculated, followed by determining the MTOC’s distance and orientation relative to the nucleus. To categorize distinct zones within the 1x dead end-channels, their x/y coordinates, and orientations are initially computed using template matching. Subsequently, data from all tracks is consolidated, and maps depicting speed, orientation, and MTOC distance are generated for in-depth analysis.

### Statistics

All replicates were validated independently and pooled only when all showed similar results. Statistical analysis was conducted using GraphPad Prism using the appropriate tests according to normal or non-normal data distribution as stated in the figure legends. Error bars are defined in the figure legends.

## Acknowledgements

We thank Christoph Mayr, Bingzhi Wang, and Artur Kuznetcov for initial experiments on amoeboid nucleokinesis, Daniela Rieger for technical assistance with *Dictyostelium* experiments, Ana-Maria Lennon-Duménil and Aline Yatim for bone marrow from MyoIIA-Flox*CD11c-Cre mice, Michael Sixt and Aglaja Kopf for EMTB-mCherry, EB3-mCherry, and Myh9-GFP expressing HoxB8 cells, Malte Benjamin Braun, Mauricio Ruiz, and Madeleine T. Schmitt for critical reading of the manuscript, and the Core Facility Bioimaging, the Core Facility Flow Cytometry, and the Animal Core Facility of the Biomedical Center (BMC) for excellent support. This study was supported by the Peter Hans Hofschneider Professorship of the foundation ‘Stiftung Experimentelle Biomedizin’, the LMU Institutional Strategy LMU-Excellent within the framework of the German Excellence Initiative, and the Deutsche Forschungsgemeinschaft (DFG; German Research Foundation; SFB914 project A12).

## Author contributions

Conceptualization: JK, JR

Investigation: JK, KS, MDH, JR

Methodology: JK, JM, LS

Software: RH

Resources: AMT

Funding acquisition: JR

Project administration: JR

Supervision: JR

Writing – original draft: JK, JR

Writing – review & editing: JK, RH, KS, MDH, JM, LS, AMT, JR

## Data Availability

This study includes no data deposited in external repositories.

## Expanded View Figure Legends

**Expanded View Figure 1.**
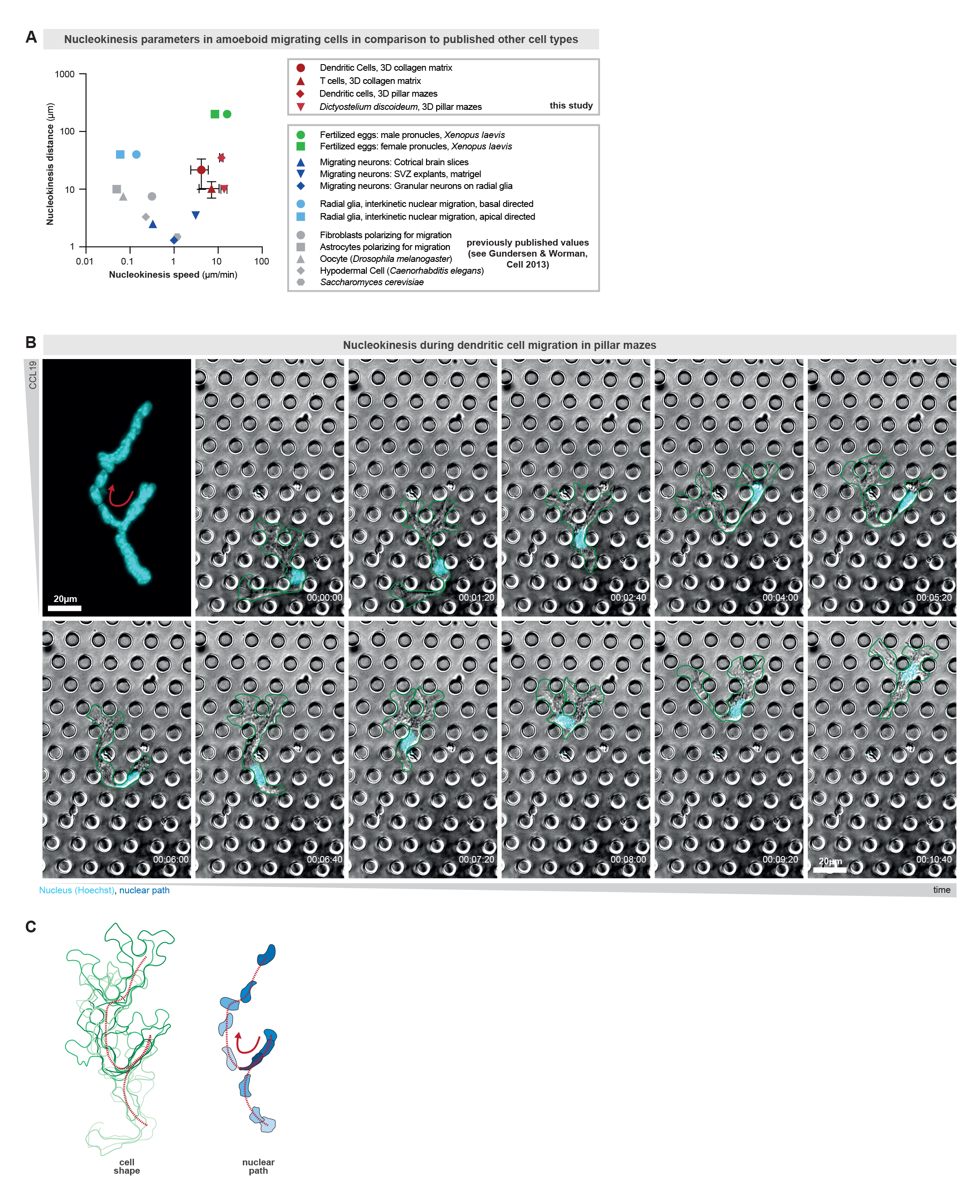
Comparison of amoeboid nucleokinesis parameters to other forms of nucleokinesis. **A.** Speed and distance of amoeboid nucleokinesis in dendritic cells, T cells, and *Dictyostelium discoideum*. Previously published nucleokinesis speeds and distances measured in other cell types are shown as a reference (Gundersen & Worman, Cell 2013). Data from this study are mean ±SD. **B.** Representative bone marrow-derived dendritic cell (DC) migrating through a porous maze-like microenvironment composed of pillars interconnecting two surfaces below and above the migrating cell. The cell shape is outlined in green and the nucleus is visualized by Hoechst (cyan). Red arrow highlights nucleokinesis event. Time in hr:min:sec. **C.** Time projections of the cellular and nuclear paths of the cell in B are shown in shades of green and blue, respectively. Red arrow highlights nucleokinesis event, and the red dashed line shows the nuclear path.

**Expanded View Figure 2.**
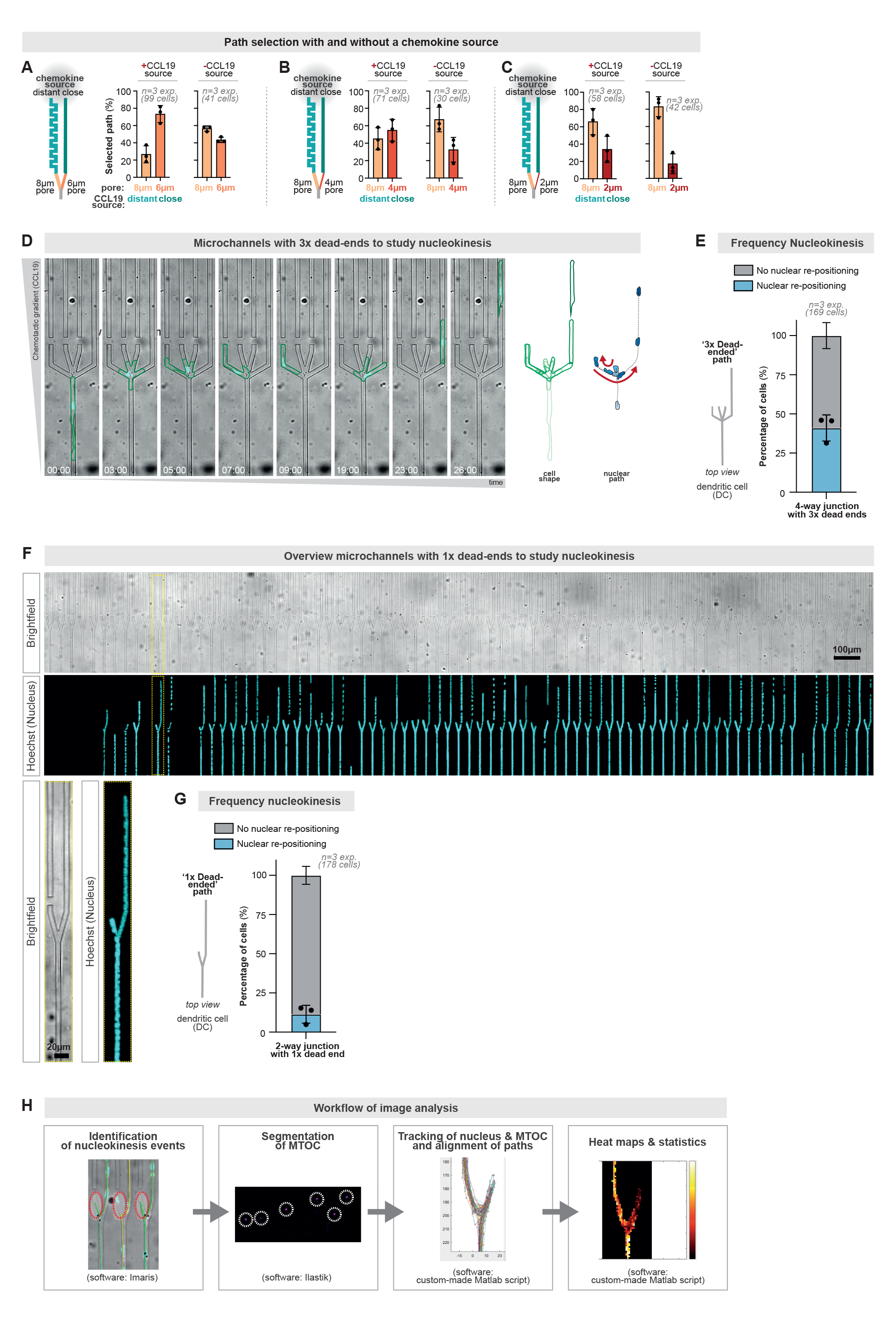
Novel reductionistic assays to characterize nucleokinesis during cell motility. **A.** Quantification of cellular path decisions in the microenvironments shown in Fig 2A with and without chemokines source. N=3 replicates (99 cells +CCL19; 41 cells without CCL19). Note that the comparative dataset of cells with CCL19 is the same data as in main Fig 2D. Data are Mean±SD. **B.** Quantification of cellular path decisions in the microenvironments shown in Fig 2B with and without chemokines source. N=3 replicates (71 cells +CCL19; 30 cells without CCL19). Note that the comparative dataset of cells with CCL19 is the same data as in main Fig 2E. Data are Mean±SD. **C.** Quantification of cellular path decisions in the microenvironments shown in Fig 2C with and without chemokines source. N=3 replicates (58 cells +CCL19; 42 cells without CCL19). Note that the comparative dataset of cells with CCL19 is the same data as in main Fig 2F. Data are Mean±SD. **D.** Representative bone marrow-derived DC (BMDC) migrating in a microchannel with a path junction that has one open and three blocked paths. Note the high frequency of nucleokinesis (nucleus in cyan; Hoechst) as quantified in E). To quantitively analyze the detailed spatio-temporal dynamics of nucleokinesis, we analyzed nucleokinesis in even more simple environments, in which cells approach path junctions that are composed of one dead-end path and one open continuous path (see F and G). The cellular shape and the position of the nucleus are outlined over time in shades of green and blue, respectively. Time in min:sec. **E.** Quantification of the rate of nucleokinesis in a 4-way junction with 3x dead ends. N=3 replicates, 169 cells. Data are mean ±SD. **F.** Overview of microchannels with 1x dead-end. Top: Brightfield, bottom: maximum projection of nuclear channel (Hoechst, cyan). The yellow-dotted regions are shown enlarged below. **G.** Quantification of the rate of nucleokinesis in a 2-way junction with 1x dead end. N=3 replicates, 178 cells. Data are mean ±SD. **H.** Workflow of semi-automated image analysis for cells migrating in the 2-junction with 1x dead end. Nucleokinesis events are identified by only selecting nuclear tracks entering a surface drawn manually over the dead end in Imaris. Ilastik is used to segment the MTOC signal. Subsequently, the nucleus and MTOC are tracked in Imaris. Using a custom-made Matlab script, tracks are aligned and heat maps and data on track speed, distance of nucleus and MTOC, and distance changes are generated.

**Expanded View Figure 3.**
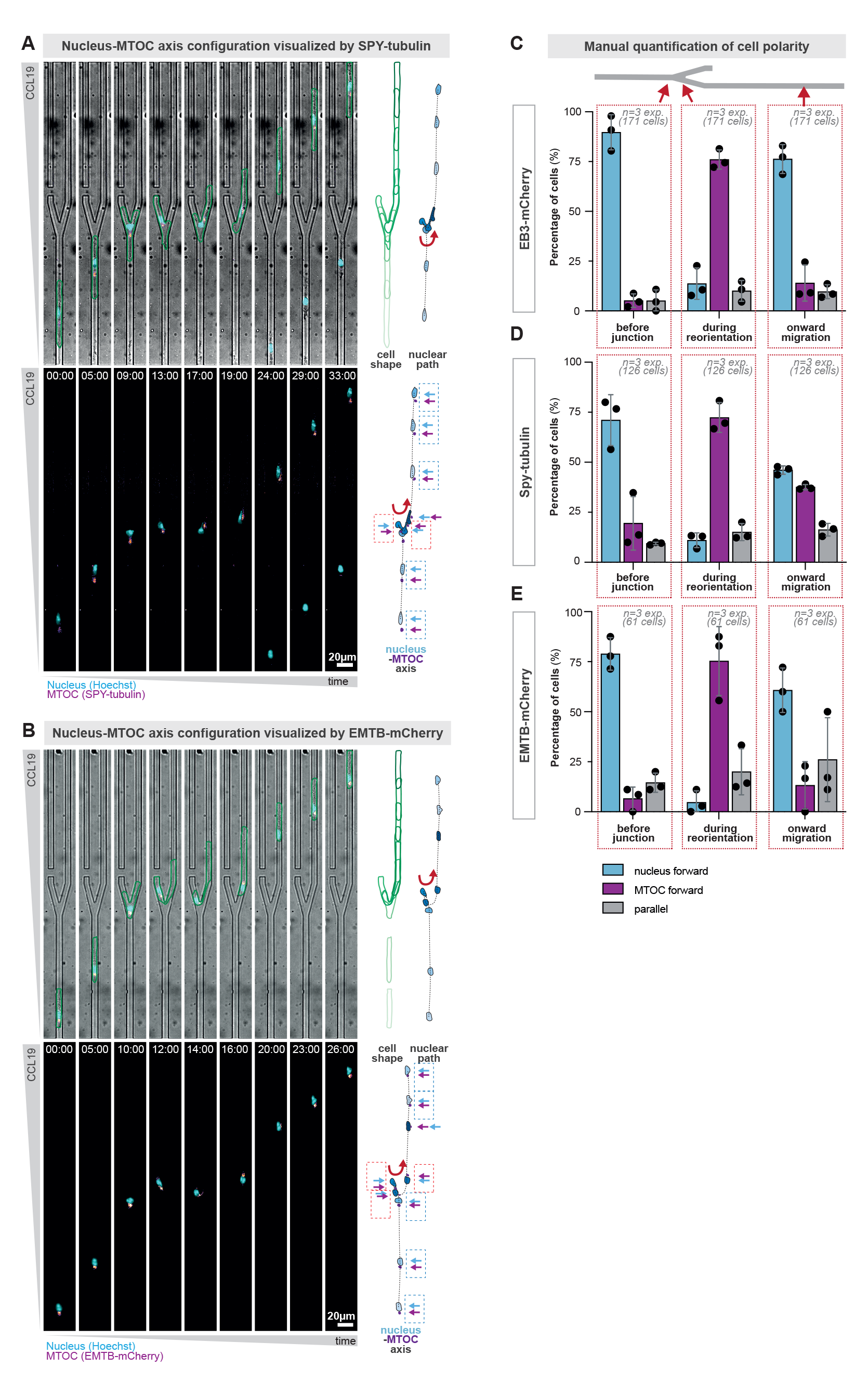
Quantification of re-orientations in the nucleus-MTOC axis during amoeboid nucleokinesis. **A.** Representative Hoxb8-derived dendritic cell (DC) approaching a path decision with equal pores sizes but one blocked path, frequently causing nucleokinesis from the blocked to the open path. The DC is stained with SPY-tubulin, which also visualizes the microtubule-organizing center (MTOC; in pink). The nucleus is visualized by Hoechst (cyan) and the cell shape is outlined in green. Projections of cellular (green), MTOC (pink) and nuclear (blue) paths are shown on the right. The configuration of the nucleus-MTOC axis is highlighted by dashed boxes (blue=nucleus forward; red=MTOC forward). Time in min:sec. **B.** As in A), but using DCs that stably encode EMTB-mCherry, which visualizes the microtubule-organizing center. **C.** Manual quantification of the nucleus-MTOC axis configuration in EB3-mCherry expressing Hoxb8-derived dendritic cells before, during, and after nucleokinesis. The nucleus-MTOC axis configuration was assessed at the channel regions indicated by the red arrows. N=3 replicates, 171 cells. Data are mean ±SD. **D.** Manual quantification of the nucleus-MTOC axis configuration in SPY-tubulin stained Hoxb8-derived dendritic cells before, during, and after nucleokinesis. The nucleus-MTOC axis configuration was assessed at the channel regions indicated by the red arrows. N=3 replicates, 126 cells. Data are mean ±SD. **E.** Manual quantification of the nucleus-MTOC axis configuration in EMTB-mCherry expressing Hoxb8-derived dendritic cells before, during, and after nucleokinesis. The nucleus-MTOC axis configuration was assessed at the channel regions indicated by the red arrows. N=3 replicates, 61 cells. Data are mean ±SD.

**Expanded View Figure 4.**
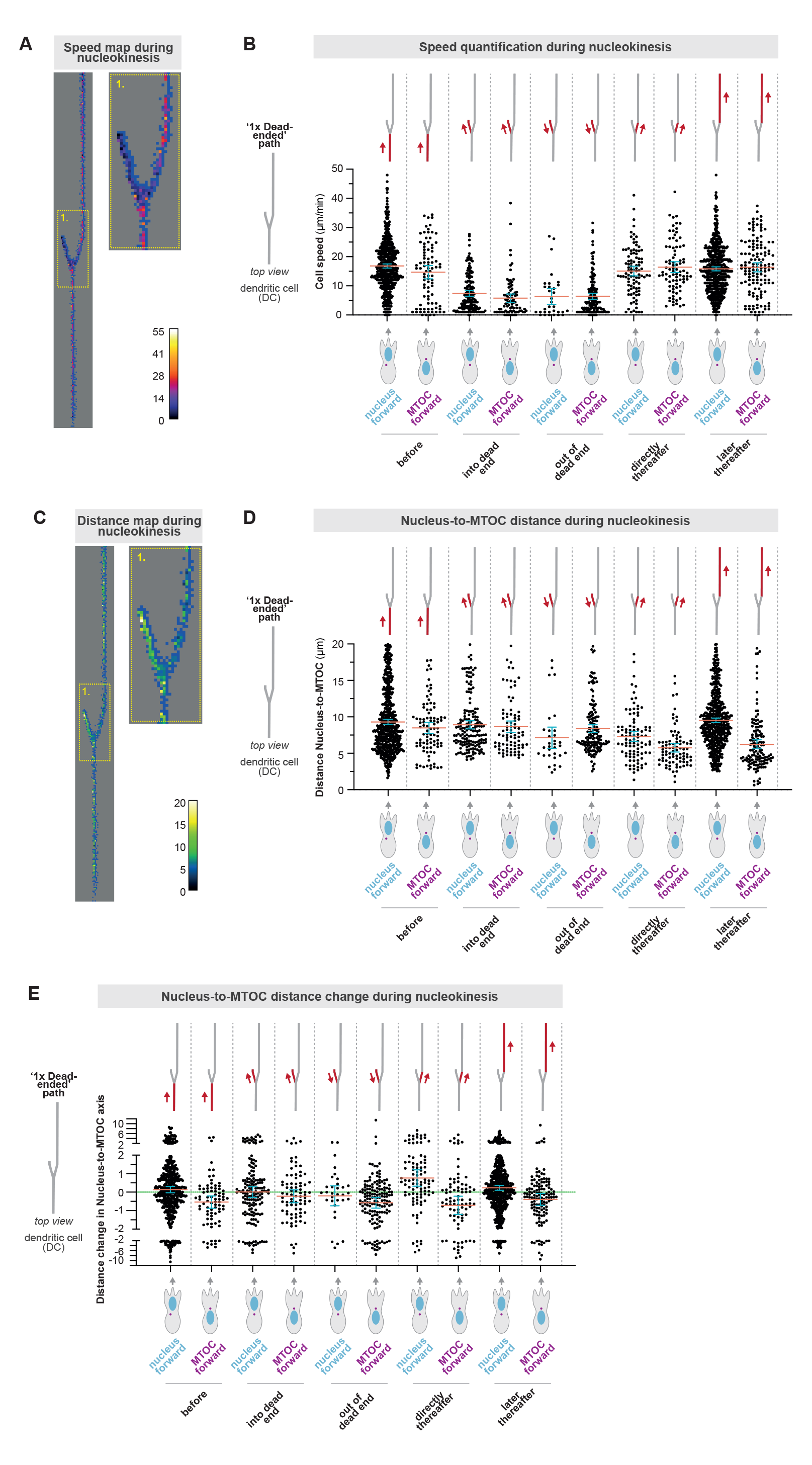
Nucleus-to-MTOC distances and speeds during amoeboid nucleokinesis. **A.** Heatmap of nuclear speed during amoeboid nucleokinesis. The yellow-dotted region 1 is enlarged to depict the cellular behavior during the initial nucleokinesis event. N=6 replicates, 48 cells. **B.** Quantification of nuclear speed during amoeboid nucleokinesis depending on the nucleus-MTOC axis configuration: EB3-mCherry expressing HoxB8-derived DCs, migrating through a path junction with one blocked path. The data show tracked nuclear velocities (visualized by Hoechst) of DCs that reposition their nucleus from the blocked to the open path by nucleokinesis. N=6 replicates, N=48 cells, total analyzed events per column (from 1^st^ to last column): 566, 90, 165, 85, 32, 159, 96, 85, 541, 136; Data are mean ±95CI. **C.** Heatmap of the distance between nucleus and MTOC during amoeboid nucleokinesis. The yellow-dotted region 1 is enlarged to depict the cellular behavior during the initial nucleokinesis event. N=6 replicates, 48 cells. **D.** Quantification of the distance between nucleus and MTOC during amoeboid nucleokinesis depending on the nucleus-MTOC axis configuration: EB3-mCherry expressing HoxB8-derived DCs, migrating through a path junction with one blocked path. The data show the distances between the center of the nucleus (visualized by Hoechst) and the center of the MTOC (visualized by EB3-mCherry) in DCs that reposition their nucleus from the blocked to the open path by nucleokinesis. N=6 replicates, N=48 cells, total analyzed events per column (from 1^st^ to last column): 566, 90, 165, 85, 32, 159, 96, 85, 541, 136; Data are mean ±95CI. **E.** Quantification of the nucleus-to-MTOC distance changes during amoeboid nucleokinesis depending on the nucleus-MTOC axis configuration: EB3-mCherry expressing HoxB8-derived DCs, migrating through a path junction with one blocked path. The data show the change of distances between the center of the nucleus (visualized by Hoechst) and the center of the MTOC (visualized by EB3-mCherry) in DCs that reposition their nucleus from the blocked to the open path by nucleokinesis. N=6 replicates, N=48 cells, total analyzed events per column (from 1^st^ to last column): 566, 90, 165, 85, 32, 159, 96, 85, 541, 136; Data are mean ±95CI.

**Expanded View Figure 5.**
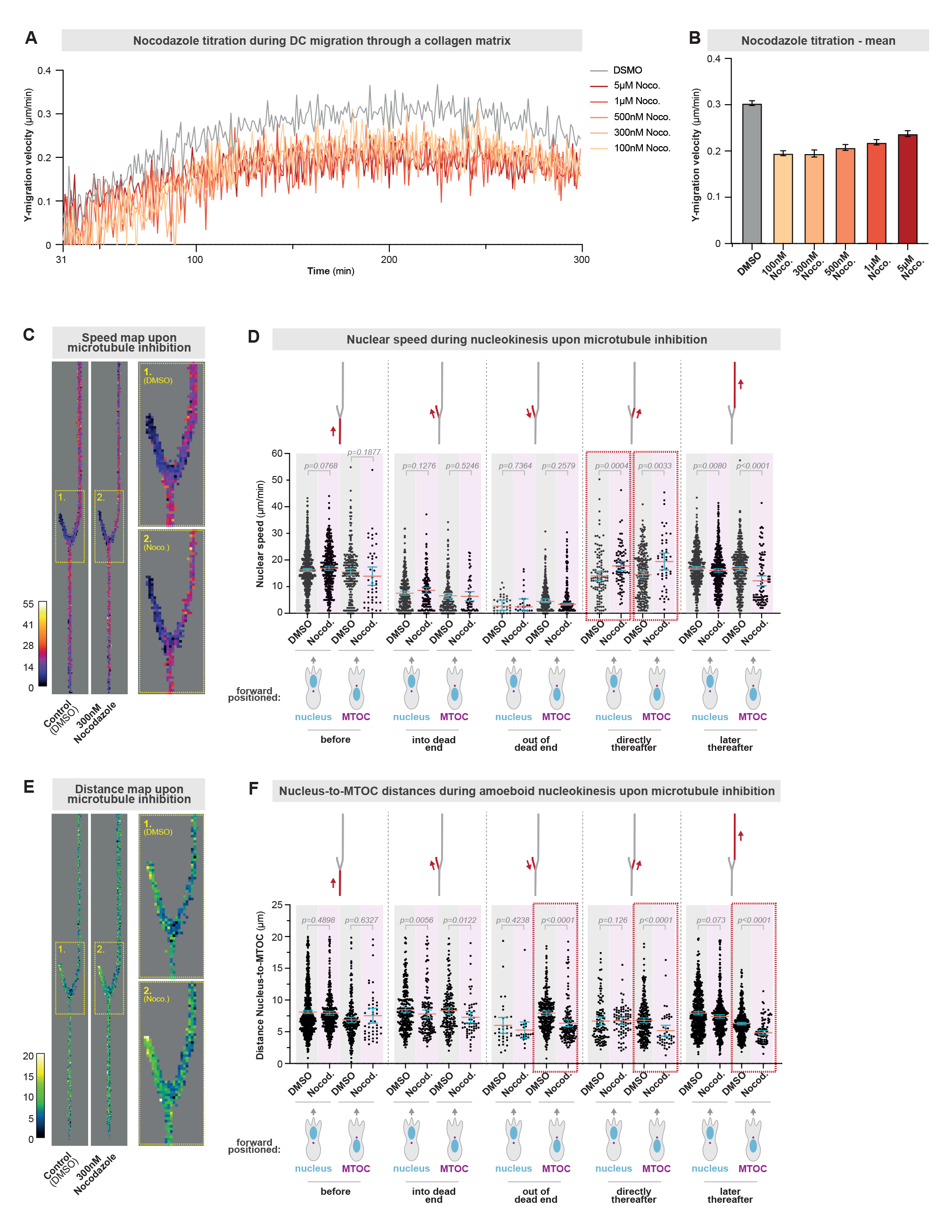
Faster amoeboid nucleokinesis when microtubules are depolymerized. **A.** Bone marrow-derived dendritic cell migration in three-dimensional (3D) collagen matrices (3.3 mg/ml) along a CCL19 chemokine gradient in the presence of different concentrations of Nocodazole (microtubule inhibitor) or DMSO (control). N=3 replicates. Data are mean. **B.** Mean migration velocity between 150 and 250 min of cells shown in A. N=3 replicates. Data are mean ±95CI. **C.** Heatmap of nuclear speed during amoeboid nucleokinesis in the presence of the microtubule inhibitor nocodazole (300 nM) or control (DMSO). The yellow-dotted regions 1 (DMSO) and 2 (Nocodazole) are enlarged to depict the cellular behavior during the initial nucleokinesis event. N=3 replicates, 96 (DMSO) and 38 (Nocodazole) cells. **D.** Quantification of nuclear speed during amoeboid nucleokinesis upon microtubule inhibition with 300nM nocodazole (Nocod.): EMTB-mCherry expressing HoxB8-derived DCs, migrating through a path junction with one blocked path. The data show tracked nuclear velocities (visualized by Hoechst) of DCs that reposition their nucleus from the blocked to the open path by nucleokinesis. The red dashed boxes highlight the accelerated nuclear speed during repositioning of the nucleus in the open path. N=3 replicates, 96 (DMSO) and 38 (Nocodazole), total analyzed events per column (from 1^st^ to last column): 859, 417, 235, 45, 305, 145, 188, 58, 34, 23, 304, 149, 111, 81, 249, 46, 577, 505, 484, 97; mean±95CI, Mann-Withney test. **E.** Heatmap of distance between nucleus and MTOC during amoeboid nucleokinesis in the presence of the microtubule inhibitor nocodazole (300 nM) or control (DMSO). The yellow-dotted regions 1 (DMSO) and 2 (Nocodazole) are enlarged to depict the cellular behavior during the initial nucleokinesis event. N=3 replicates, 96 (DMSO) and 38 (Nocodazole) cells. **F.** Quantification of the distance between nucleus and MTOC during amoeboid nucleokinesis upon microtubule inhibition with 300nM nocodazole (Nocod.): EMTB-mCherry expressing HoxB8-derived DCs, migrating through a path junction with one blocked path. The data show the distances between the center of the nucleus (visualized by Hoechst) and the center of the MTOC (visualized by EMTB-mCherry) in DCs that reposition their nucleus from the blocked to the open path by nucleokinesis. The red dashed boxes highlight the reduced distance between nucleus and MTOC upon microtubule inhibition, when the MTOC is positioned frontward. N=3 replicates, 96 (DMSO) and 38 (Nocodazole), total analyzed events per column (from 1^st^ to last column): 859, 417, 237, 46, 305, 145, 188, 58, 34, 23, 304, 149, 111, 81, 249, 46, 578, 508, 484, 97; mean±95CI; Mann-Whitney test.

**Expanded View Figure 6.**
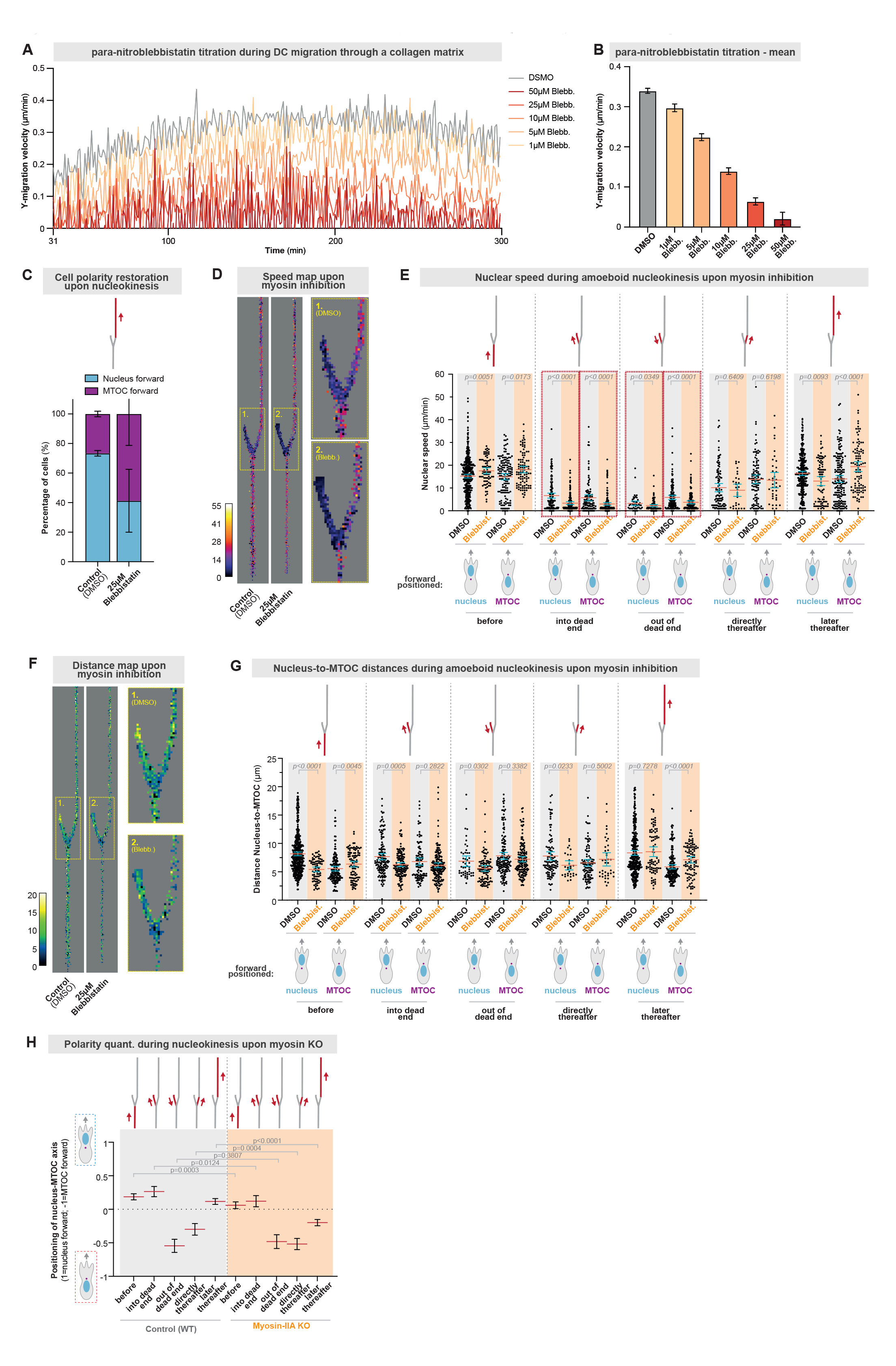
Reduced amoeboid nucleokinesis speed and cell polarity switching upon myosin inhibition. **A.** Bone marrow-derived dendritic cell migration in three-dimensional (3D) collagen matrices (3.3 mg/ml) along a CCL19 chemokine gradient in the presence of different concentrations of para-nitroblebbistatin (myosin inhibitor) or DMSO (control). N=3 replicates. Data are mean. **B.** Mean migration velocity between 150 and 250 min of cells shown in A. N=3 replicates. Data are mean ±95CI. **C.** Quantification of the cell polarity after nucleokinesis at the channel exit in the presence of the myosin inhibitor para-nitroblebbistatin (25 µM) or control (DMSO). N=4 replicates, 37 (DMSO) and 13 (para-nitroblebbistatin) cells. Data are mean ±SEM. **D.** Heatmap of nuclear speed during amoeboid nucleokinesis in the presence of the myosin inhibitor para-nitroblebbistatin (25 µM) or control (DMSO). The yellow-dotted regions 1 (DMSO) and 2 (para-nitroblebbistatin) are enlarged to depict the cellular behavior during the initial nucleokinesis event. N=4 replicates, 37 (DMSO) and 13 (Blebbist.) cells. **E.** Quantification of nuclear speed during amoeboid nucleokinesis upon myosin inhibition with 25μM para-nitroblebbistatin (Blebbist.): EB3-mCherry expressing HoxB8-derived DCs, migrating through a path junction with one blocked path. The data show tracked nuclear velocities (visualized by Hoechst) of DCs that reposition their nucleus from the blocked to the open path by nucleokinesis. The red dashed boxes highlight the decelerated nuclear speed upon myosin inhibition during repositioning of the nucleus from the blocked path. N=4 replicates, 37 (DMSO) and 13 (Blebbist.) cells; total analyzed events per column (from 1^st^ to last column): 433, 76, 146, 97, 139, 147, 107, 168, 48, 99, 150, 145, 101, 29, 107, 38, 330, 83, 184, 97; mean±95CI, Mann-Withney test. **F.** Heatmap of distance between nucleus and MTOC during amoeboid nucleokinesis in the presence of the myosin inhibitor para-nitroblebbistatin (25 µM) or control (DMSO). The yellow-dotted regions 1 (DMSO) and 2 (Blebbist.) are enlarged to depict the cellular behavior during the initial nucleokinesis event. N=4 replicates, 37 (DMSO) and 13 (Blebbist.) cells. **G.** Quantification of the distance between nucleus and MTOC during amoeboid nucleokinesis upon myosin inhibition with 25 µM para-nitroblebbistatin (Blebbist.): EB3-mCherry expressing HoxB8-derived DCs, migrating through a path junction with one blocked path. The data show the distances between the center of the nucleus (visualized by Hoechst) and the center of the MTOC (visualized by EB3-mCherry) in DCs that reposition their nucleus from the blocked to the open path by nucleokinesis. N=4 replicates, 37 (DMSO) and 13 (Blebbist.) cells, total analyzed events per column (from 1^st^ to last column): 433, 76, 146, 97, 139, 147, 107, 168, 48, 99, 150, 145, 101, 29, 107, 38, 330, 83, 184, 97; mean±95CI, Mann-Withney test. **H.** Quantification of the nucleus-MTOC axis configuration before, during, directly after, and later after amoeboid nucleokinesis upon myosin-IIA KO or control (1=all cells position the nucleus in front of the MTOC; −1= all cells position the MTOC in front of the nucleus). Data are mean±95CI, Mann-Whitney test, N=3 replicates, 119 (control) and 94 (myosin KO) cells, and 1813 (before; control), 1511 (before; KO), 617 (into a dead end, control), 555 (into a dead end, KO), 286 (out of a dead end, control), 282 (out of a dead end, KO), 468 (directly thereafter, control), 411 (directly thereafter, KO), 1953 (later thereafter, control), and 1700 (later thereafter, KO) image frames.

**Expanded View Figure 7.**
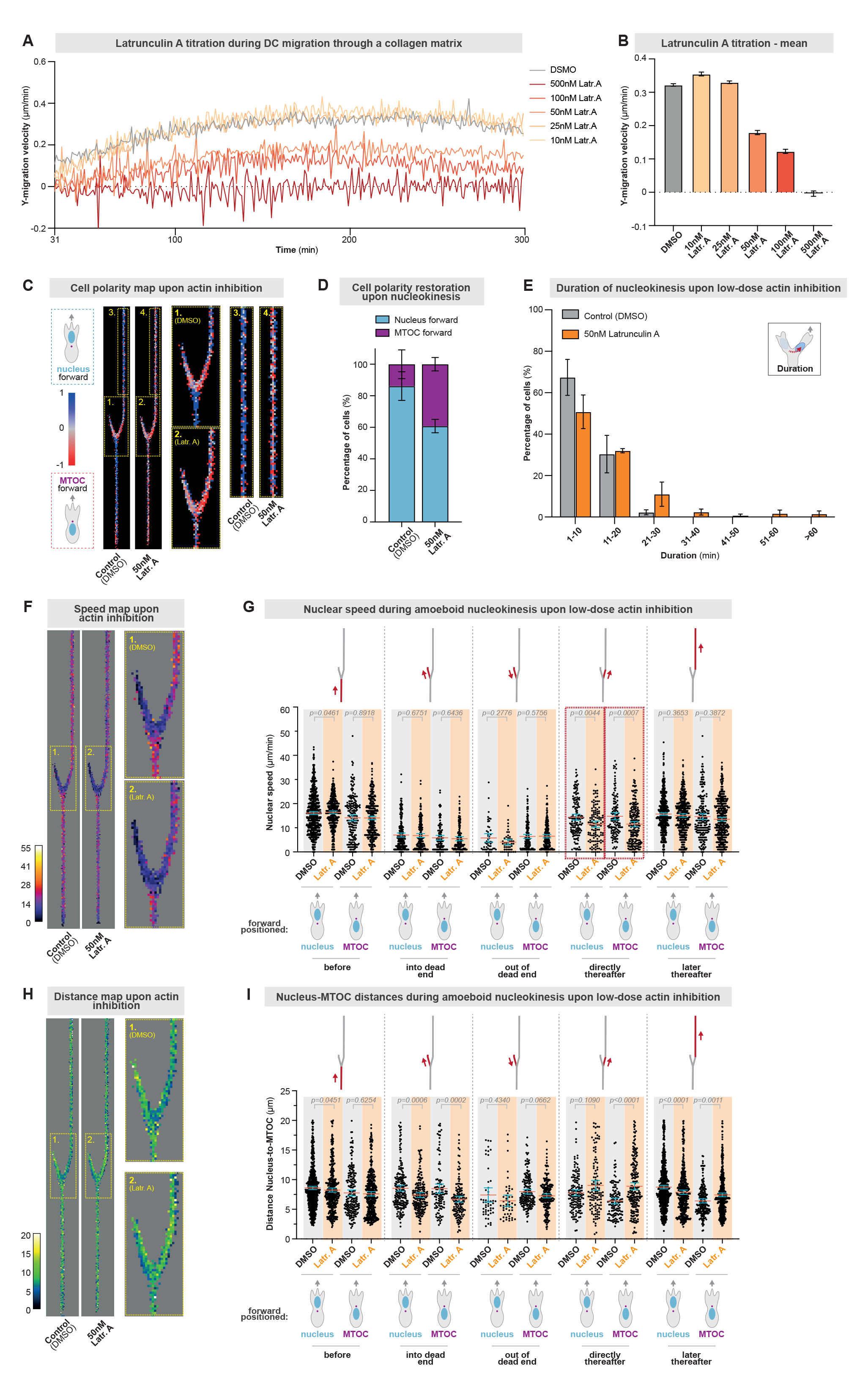
Delayed cell polarity switching and amoeboid nucleokinesis upon low-dose actin inhibition. **A.** Bone marrow-derived dendritic cell migration in three-dimensional (3D) collagen matrices (3.3 mg/ml) along a CCL19 chemokine gradient in the presence of different concentrations of Latrunculin A (actin inhibitor) or DMSO (control). N=3 replicates. Data are mean. **B.** Mean migration velocity between 150 and 250 min of cells shown in A. N=3 replicates. Data are mean ±95CI. **C.** Heatmap of the nucleus-MTOC axis configuration during amoeboid nucleokinesis in the presence of the actin inhibitor Latrunculin A (50 nM) or control (DMSO). The yellow-dotted regions 1 (DMSO) and 2 (Latrunculin A) are enlarged to depict the cellular behavior during the initial nucleokinesis event, and the yellow-dotted regions 3 (DMSO) and 4 (Latrunculin A) are enlarged to depict the cellular behavior during the later nucleokinesis events to reposition the nucleus to the cellular front. N=3 replicates, 78 (DMSO) and 67 (Latrunculin A) cells. **D.** Quantification of the cell polarity after nucleokinesis at the channel exit in the presence of the actin inhibitor Latrunculin A (50 nM) or control (DMSO). N=3 replicates, 78 (DMSO) and 66 (Latrunculin A) cells. Data are mean ±SEM. **E.** Quantification of the duration of nucleokinesis in the presence of the actin inhibitor Latrunculin A (50 nM) or control (DMSO). N=3 replicates, 101 (DMSO) and 93 (Latrunculin A) cells. Data are mean ±SEM. **F.** Heatmap of nuclear speed during amoeboid nucleokinesis in the presence of the actin inhibitor Latrunculin A (50 nM) or control (DMSO). The yellow-dotted regions 1 (DMSO) and 2 (Latrunculin A) are enlarged to depict the cellular behavior during the initial nucleokinesis event. N=3 replicates, 78 (DMSO) and 67 (Latrunculin A) cells. **G.** Quantification of nuclear speed during amoeboid nucleokinesis upon actin inhibition with 50 nM Latrunculin A (Latr. A): EB3-mCherry expressing HoxB8-derived DCs, migrating through a path junction with one blocked path. The data show tracked nuclear velocities (visualized by Hoechst) of DCs that reposition their nucleus from the blocked to the open path by nucleokinesis. The red dashed boxes highlight the decelerated nuclear speed upon actin inhibition during repositioning of the nucleus from the blocked path. N=3 replicates, 78 (DMSO) and 67 (Latr. A) cells, total analyzed events per column (from 1^st^ to last column): 684, 444, 206, 413, 210, 204, 113, 147, 50, 48, 184, 217, 155, 119, 127, 209, 700, 434, 201, 465; data are mean±95CI, Mann-Withney test. **H.** Heatmap of distance between nucleus and MTOC during amoeboid nucleokinesis in the presence of the actin inhibitor Latrunculin A (50 nM) or control (DMSO). The yellow-dotted regions 1 (DMSO) and 2 (Latr. A) are enlarged to depict the cellular behavior during the initial nucleokinesis event. N=3 replicates, 78 (DMSO) and 67 (Latr. A) cells. **I.** Quantification of the distance between nucleus and MTOC during amoeboid nucleokinesis upon actin inhibition with 50 nM Latrunculin A (Latr. A): EB3-mCherry expressing HoxB8-derived DCs, migrating through a path junction with one blocked path. The data show the distances between the center of the nucleus (visualized by Hoechst) and the center of the MTOC (visualized by EB3-mCherry) in DCs that reposition their nucleus from the blocked to the open path by nucleokinesis. N=3 replicates, 78 (DMSO) and 67 (Latr. A) cells, total analyzed events per column (from 1^st^ to last column): 684, 444, 206, 413, 210, 204, 113, 147, 50, 48, 184, 217, 155, 119, 127, 209, 700, 434, 201, 465; data are mean±95CI, Mann-Withney test.

**Expanded View Figure 8.**
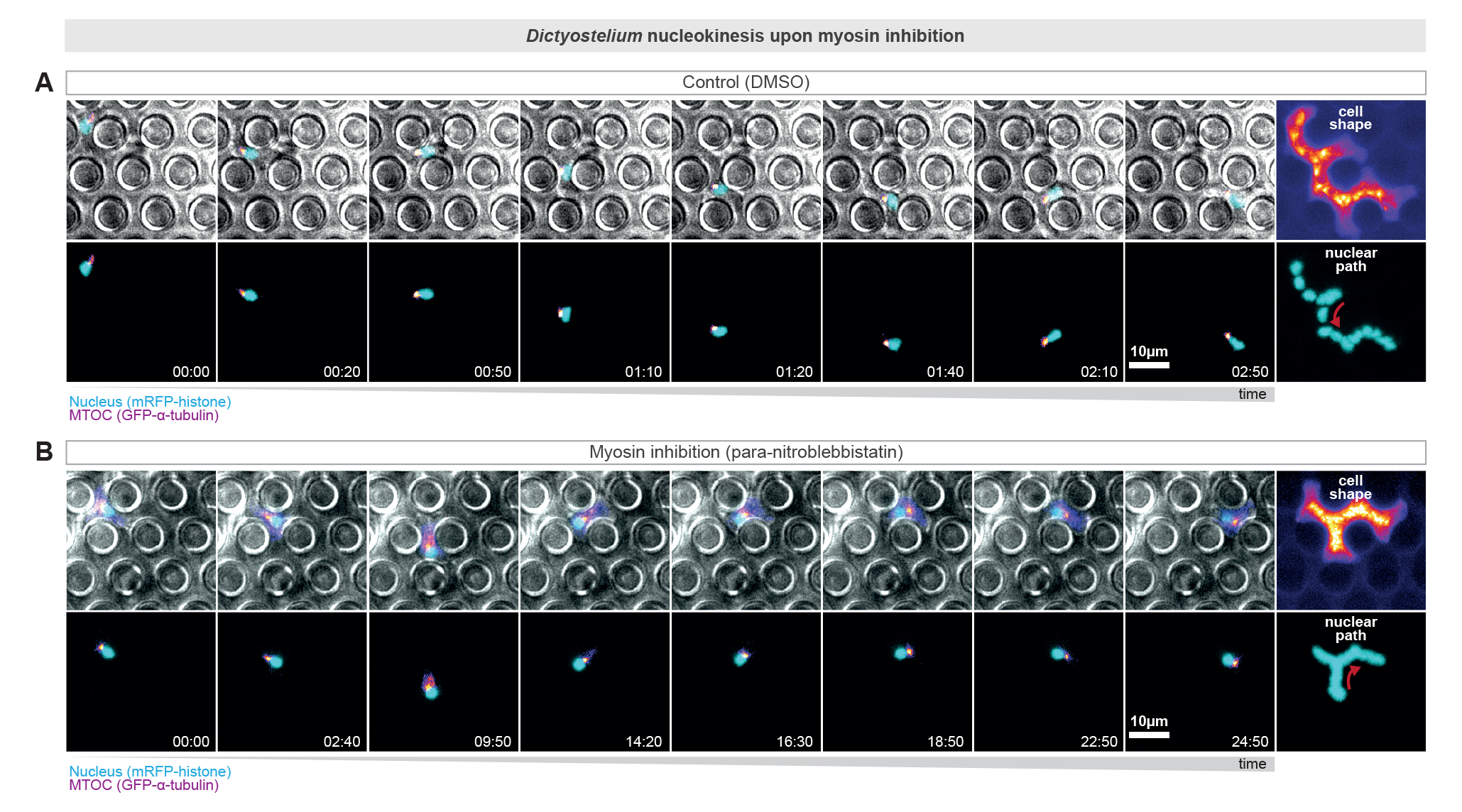
Delayed nucleokinesis upon myosin inhibition in *Dictyostelium*. **A.** Representative *Dictyostelium* cell (control: DMSO) migrating in a pillar maze. The *Dictyostelium* cell stably encodes GFP-α-tubulin, which also visualizes the microtubule-organizing center (MTOC; in pink) and mRFP-histone, which visualizes the nucleus (in cyan). Projections of cellular (fire) and nuclear (blue) paths are shown on the right. The red arrow highlights the nucleokinesis event. Time in min:sec. **B.** Representative *Dictyostelium* cell migrating in a pillar maze upon myosin inhibition (25 µM para-nitroblebbistatin). The *Dictyostelium* cell stably encodes GFP-α-tubulin, which also visualizes the microtubule-organizing center (MTOC; in pink) and mRFP-histone, which visualizes the nucleus (in cyan). Projections of cellular (fire) and nuclear (blue) paths are shown on the right. The red arrow highlights the nucleokinesis event. Time in min:sec.

## Expanded View Movies

**Expanded View Movie 1.**

Nucleokinesis during amoeboid immune cell migration in three-dimensional collagen networks. Representative dendritic cell (1^st^ movie part) and T cell (2^nd^ movie part) migrating in three-dimensional collagen networks along a CCL19 chemokine gradient. The nucleus is shown in cyan (Hoechst), and full and partial nucleokinesis events are highlighted by red arrows. The movie shows representative cells from at least three independent biological replicates. Time is indicated as min:sec.

**Expanded View Movie 2.**

Adaptive pathfinding in microenvironments of competing chemokine and pore size cues. Representative dendritic cells (DCs) stained with Hoechst (nuclear label; cyan) passing a path decision junction, at which migrating cells encounter two path options with different strengths of chemotactic as well as pore size cues. The first movie part shows DCs in which the cellular and nuclear paths were identical (left panel: 8- vs. 6-micrometer pores; middle panel: 8- vs. 4-micrometer pores; right panel: 8- vs. 2-micrometer pores. The left path is always more distant and the right path is closer to the CCL19 chemokine source). The path chosen by each cell is highlighted by red arrows. The second movie part shows representative examples of full (left panel) and partial (right panel) nuclear repositioning events to adapt the nuclear path to the cellular path. Full and partial nucleokinesis events are highlighted by red arrows. The movie shows representative cells from at least three independent biological replicates. Time is indicated as min:sec.

**Expanded View Movie 3.**

Polarity of the nucleus-to-MTOC axis configuration during amoeboid nucleokinesis. Representative EB3-mCherry expressing dendritic cell (first movie part), EMTB-mCherry expressing dendritic cell (second movie part), and Spy-tubulin labeled dendritic cell (third movie part) migrating through a path junction. The nucleus is labeled with Hoechst (cyan) and EB3-mCherry/EMTB-mCherry/Spy-tubulin signals are shown in fire color-code. The movie shows representative cells from at least three independent biological replicates. Time is indicated as min:sec.

**Expanded View Movie 4.**

Amoeboid nucleokinesis in microtubule-inhibited cells. Representative dendritic cells (DCs) with a nuclear label (Hoechst; cyan) and a MTOC label (EMTB-mCherry; fire color-coded) migrating through a path junction and performing nucleokinesis. The left movie part shows a control DC (DMSO) and the right part a microtubule-inhibited DC (300nM Nocodazole; right panel). The movie shows representative cells from at least three independent biological replicates. Time is indicated as min:sec.

**Expanded View Movie 5.**

Myosin localization during amoeboid nucleokinesis. Representative Myh9-GFP (encoding Myosin-IIA-GFP; fire-color coded) expressing dendritic cell migrating in pillar mazes. The nucleus is labeled with Hoechst (cyan). Nuclear repositioning is highlighted by a white arrow. The movie shows representative cells from at least three independent biological replicates. Time is indicated as min:sec.

**Expanded View Movie 6.**

Amoeboid nucleokinesis in myosin-inhibited cells. Representative dendritic cells with a nuclear label (Hoechst; cyan) and a MTOC label (EB3-mCherry; fire color-coded) migrating through a path junction and performing nucleokinesis. The movie shows myosin-inhibited (para-nitroblebbistatin; right panel) and control (DMSO; left panel) cells, representative for at least three independent biological replicates. Time is indicated as h:min:s.

**Expanded View Movie 7.**

Nucleokinesis during *Dictyostelium discoideum* and dendritic cell migration. Representative dendritic cell (1^st^ movie part) and *Dictyostelium discoideum* cell (2^nd^ movie part) migrating in a pillar maze and performing nucleokinesis. The nucleus of the dendritic cell is labeled with Hoechst (cyan), and the *Dictyostelium* cell stably expresses mRFP-histone (nuclear marker; cyan). Nucleokinesis events are highlighted by red arrows. The movie shows representative cells from at least three independent biological replicates. Time is indicated as min:sec.

**Expanded View Movie 8.**

Polarity of the nucleus-to-MTOC axis configuration during nucleokinesis in *Dictyostelium discoideum*. Representative mRFP-histone (nuclear marker; cyan) and GFP-α-tubulin (MTOC marker; fire color-coded) expressing *Dictyostelium* cells. Nuclear repositioning is highlighted by a red arrow. The movie shows representative cells from at least three independent biological replicates. Time is indicated as min:sec.

**Expanded View Movie 9.**

Amoeboid nucleokinesis in *Dictyostelium discoideum* upon myosin inhibition or myosin-mutant expression. Representative *Dictyostelium* cells with a nuclear label (mRFP-histone; cyan) and a MTOC label (GFP-α-tubulin; fire color-coded) migrating through path junctions and performing nucleokinesis. The first movie part shows wild-type myosin-inhibited (para-nitroblebbistatin; bottom panel) and control (DMSO; top panel) cells, the second movie part shows myosin mutant expressing (right panel) and control (left panel) cells. Nucleokinesis events are highlighted by red arrows. The movie shows representative cells from at least three independent biological replicates. Time is indicated as min:sec.

